# Neighbor density-dependent facilitation promotes coexistence and internal oscillation

**DOI:** 10.1101/2025.02.11.636978

**Authors:** Lisa Buche, Lauren G. Shoemaker, Lauren M. Hallett, Peter Vesk, Oscar Godoy, Margaret Mayfield

## Abstract

The ability of species to form diverse communities is not fully understood. Species are known to interact in various ways with their neighborhood. Despite this, common models of species coexistence assume that per capita interactions are constant and competitive, even as the environment changes. In this study, we investigate how neighbor density-dependent variation in the strength and sign of species interactions changes species and community dynamics. We show that by including these sources of variation, predictions of ecological dynamics are significantly improved compared to outcomes of typical models that hold interaction strengths constant. We compared how well models based on different functions of neighbor density and identity did in describing population trajectories (i.e., persistence over time) and community dynamics (i.e., temporal stability, synchrony and degree of oscillation) in simulated two-species communities and a real diverse annual plant system. In our simulated communities, we found the highest level of coexistence between species pairs when species interactions varied from competitive to facilitative according to neighbor density (i.e., following a sigmoid function). Introducing within-guild facilitation through a nonlinear bounded function allowed populations, both simulated and empirical, to avoid extinction or runaway growth. In fact, nonlinear bounded functions (i.e., exponential and sigmoid functions) predicted population trends over time within the range of abundances observed over the last 10 years. With the sigmoid function, the simulated communities of two species displayed a higher probability of synchrony and oscillation than other functional forms. These simulated communities did not always show temporal stability but were predicted to coexist. Overall, varying species interactions lead to realistic ecological trajectories and community dynamics when bounded by asymptotes based on neighbor density. These findings are important for advancing our understanding of how diverse communities are sustained and for operationalizing ecological theory in the study of the real world.

## 1. Introduction

A central goal of theoretical ecology is to understand how species interactions support coexistence (Levin, 1970; Vellend, 2010; Cadotte and Tucker, 2017). A classic rule for coexistence is that a species must compete with itself more than it is limited by its heterospecific neighbors (Chesson, 2000). In keeping with this rule, both theoretical and empirical tests of species coexistence focus on direct competition despite overwhelming empirical evidence that species interactions can be facilitative (Rietkerk et al., 1997; Sans et al., 1998; Bruno et al., 2003; Brooker et al., 2008; McIntire and Fajardo, 2014) and non-linear (Kéfi et al., 2012; Mayfield and Stouffer, 2017). This bias partly arises from the tendency for theory to build on previous models, creating a ‘priority effect’ for theoretical frameworks (Simha et al., 2022; Stouffer, 2022). We hypothesize that reliance on competition-based models limits our ability to accurately predict local diversity and patterns of coexistence and population growth (e.g., Matias et al. 2018; Bimler et al. 2018; Godoy 2019; Maynard et al. 2019; Rinella et al. 2020).

Interactions can switch from beneficial to harmful depending on the species identity, density, life-stage, and neighborhood heterogeneity (Callaway and Walker, 1997; Schiffers and Tielbörger, 2006; Fayolle et al., 2009). A forb species can benefit from annual grass that provides shade (see fig.1), which helps it avoid drying out and enhances its growth and flower production. However, if the number of annual grasses increases due to higher nutrient levels, the effects on the forb may turn negative (as shown in fig.1). This can happen through competition for space, increased water stress, excessive shading, or reduced visibility to pollinators (Ashman et al., 2004; Sheley and James, 2014). This switch in the strength and sign of species interactions from facilitation to competition according to neighborhood conditions can affect the ability of species to coexist (Bimler et al., 2018; Matias et al., 2018; Wainwright et al., 2019). To rigorously evaluate this possibility, we need to incorporate more complexity into our theoretical models.

**Figure 1:**
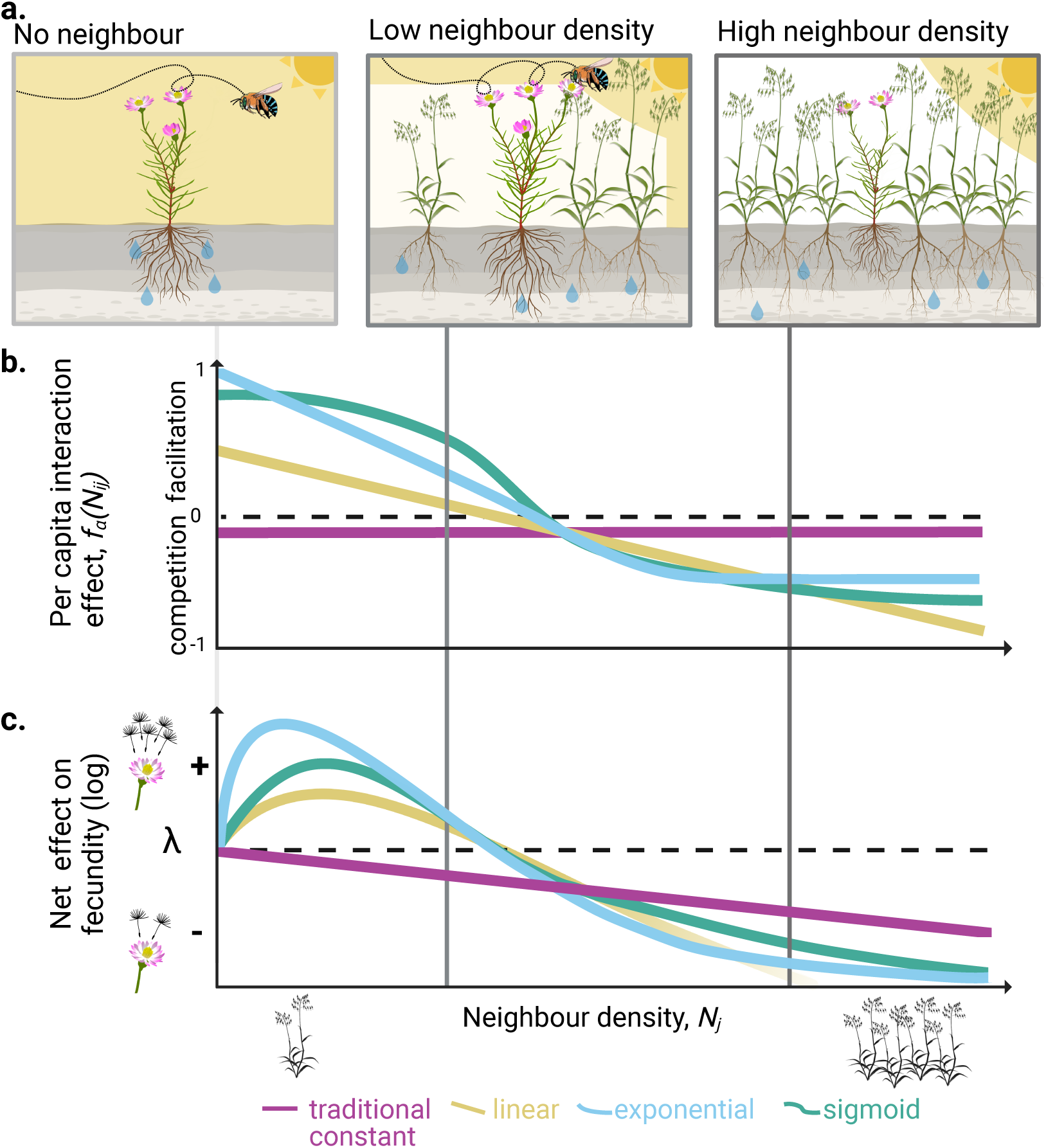
Species interactions depend on neighborhood density and identity. Take, for example, a native daisy living in an invaded grassland. Hypothetically, this daisy may compete with grasses for water, while profiting from grass-created shade. The effect of the grass on the daisy’s performance (growth, fecundity) thus depends on the number of intersecting factors involving grass individuals. **Panel a** - Shows how these interactions combine to impact the focal daisy. **Panel b.** Shows the four functional forms of interactions examined in this study as they relate to the processes shown in panel a. Each function (shown by the colored lines) has different responses to neighbor grass density. The red line shows the ‘traditional’ constant function that does not vary with neighborhood density. Yellow, blue and green lines show the three alternative functions that we propose for consideration in this study. The yellow line shows the linear function, the blue the exponential function, and the green the sigmoid function. We make the assump8tion that interactions can only decrease in all cases, going from facilitation or low competition to relatively higher competition with increasing neighbor density, as at the extreme, individuals have to compete for space. **c.** Shows the net effect of each functional form on the fecundity of *i* –*F*_*i*_–, when the intrinsic fitness –γ_*i*_– equals zero. Created with BioRender

Theoretical models are simplifications of this dizzying complexity of processes that underlie natural communities (Vellend, 2010). Though it is necessary to simplify models to some extent, oversimplification results in a lack of realism in both theory and empirical tests (but see Scher et al. 2024). For instance, models such as Lotka-Volterra (Lotka, 1925; Volterra, 1926), Beverton Holt (Watkinson, 1980), and Ricker (Ricker, 1954)) models, which collectively form the backbone of most tests of species coexistence (e.g., Lyu and Alexander (2023) and Daniel et al. (2024)), all assume that interactions between species are pairwise, direct and static. In practice, these models are implemented using the assumption that interactions are all competitive, with facilitative interactions generally restricted by bounding the values of competition coefficients (e.g., Narwani et al., 2013; Godoy and Levine, 2014; Blackford et al., 2020). Attempts to incorporate facilitation have usually treated it as a constant effect, which makes models prone to runaway growth (Martorell and Freckleton, 2014; Bimler et al., 2018; Hart, 2023); indeed, a major hurdle to incorporating facilitation is the belief that it invariably leads to unrestricted population growth (Hart, 2023). Expanding models to include facilitation requires expanding them to allow for non-linear variation in species interactions.

It is only in the last few years that any studies have shown the importance of non-linear variation to species interactions and resulting patterns of diversity (Koffel et al., 2021; Leibold et al., 2022; Spaak et al., 2023; Gómez et al., 2023). The inclusion of such variation is, however, still heavily biased towards studies of gradients of abiotic stress (Bimler et al., 2018; Raymundo et al., 2021; Reynaert et al., 2023; Danet et al., 2024). When non-linear effects are included in pairwise models of interactions, it has improved the explanation and prediction of population persistence in natural communities (Mayfield and Stouffer, 2017; Barbosa et al., 2023; Lai et al., 2024), suggesting that further research on the functional form and mechanistic underpinnings of non-linear interactions is warranted (Singh and Baruah, 2019; Bimler and Mayfield, 2023).

Common theoretical models of species coexistence ignore non-linear and facilitative interactions, potentially impacting our prediction of species persistence. While a few solutions exist to evaluate population persistence and coexistence of species when facilitation occurs (Koffel et al., 2021; Spaak et al., 2021; Allen-Perkins et al., 2023), none allow for variation in species interactions. A common metric for determining coexistence is ‘growth rate when rare’ where species can persist if they can increase from low density, and coexistence occurs when all species in a community have positive growth rates when rare (Barabás et al., 2018; Clark et al., 2024). Estimating species interactions as single competitive scalars along neighborhood density, rather than allowing for positive and/or non-linear interactions, reduces the potential for species interactions to increase from low density. For instance, following the above example, when the effect of the grass on the forb is dependent on grass density, the species’ interactions lead to the prediction of coexistence as the grass facilitates the forb at low density, preventing its desiccation (fig.1, linear, exponential and sigmoid function). Such a result is impossible if the effect of the grass on the forb is only estimated at a high density when they compete for resources. This typical practice ignores the possibility of facilitation via microenvironment modification, which only occurs at low densities. The growth rate/performance of a species at low density is influenced by different trade-offs of limiting factors compared to interaction at high density. Ignoring the natural variations in sign and strength of species interactions along a density gradient could be partially responsible for the mismatch between observations and predictions of local coexistence in real diverse systems (e.g., Matias et al. 2018; Bimler et al. 2018; Godoy 2019; Maynard et al. 2019; Rinella et al. 2020).

Beyond incorrect predictions of population persistence and community diversity, a lack of models that can accommodate a continuous change from facilitation at low density to competition at high density can also prevent us from obtaining a clear understanding of the internal and external processes driving species’ temporal stability. Traditional models of species coexistence show strictly competitive populations quickly settling on an equilibrium or subject to exponential growth when facilitation is included. However, this result deviates from most empirical evidence showing synchronous responses and ‘oscillations’, which are defined as a moving average in population size over time (also called “fluctuation”) (Zhao et al., 2020; Shoemaker et al., 2022). Discrepancies between theoretical predictions and observations are usually attributed to environmentally driven fluctuations (Tredennick et al., 2017; Firkowski et al., 2022). However, internal local processes are rarely considered to be potential drivers of species’ temporal fluctuations (Abbott, 2011). Synchrony is particularly likely when facilitation from other species only happens at low densities. Similarly, non-linear changes in species interactions can lead to cyclic oscillations of a community (Benincà et al., 2015; Stouffer et al., 2018). Thus, the non-linear variation along a positive-to-negative continuum based on neighbors’ density could potentially prevent runaway dynamics due to facilitation and yield internally-driven community dynamics compatible with what we observed in nature.

In this study, we have three goals. First, we define three functional forms accounting for non-linear effects of neighbor density along a continuum of facilitative to competitive interactions. Second, we assessed how inserting these functional forms in a plant population model impacts predictions of individual performance, population trajectory, and community compositional dynamics. Third, we directly assess whether the population ‘synchrony’ and ‘oscillations’ commonly observed in nature (Houlahan et al., 2007; Vasseur et al., 2014; Barraquand et al., 2022) can be recreated through population models that allow for variation in interaction outcomes. To achieve these goals, we use both simulated two-species communities and empirical data from an annual plant community.

## 2. Methods

### 2.1 Estimation of species interactions

#### 2.1.1 Traditional estimation of species interactions as constant

Models of population and multispecies dynamics usually evaluate changes in species densities (*N*_*i*,t+1_/*N*_*i*,t_, where *N* indicates the population size of species *i* at time t) as a function of species interactions and the intrinsic density-independent growth rate of the species (λ_*i*_), that is, individual performance without neighbors. The species interaction between species *i* and *j* is estimated by a coefficient (i.e, α_0,*i* *j*_) defining the effect of one individual of species *j* on the intrinsic performance of species *i* (Lotka, 1922). The interaction coefficients are traditionally estimated as a constant effect. For example, in the classic Ricker model (Ricker, 1954):

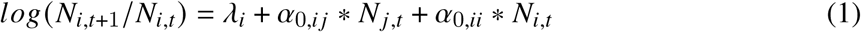

with α_0,*i* *j*_ the traditional constant inter-specific effect of one individual of species *j* on *i*, independent from neighbor density, and with α_0,*ii*_ denoting intra-specific interactions. α_0,*i* *j*_ can range from facilitative when > 0 to competitive when < 0, yet it is traditionally considered as competitive in many models to avoid runaway unbounded population growth (Hart, 2023). The interaction coefficient, which we refer to later as the net interaction outcome, is multiplied by the number of individuals of the interactive species in the community. As such, intra- and inter-specific interactions are called density-dependent, as the effect increases with the number of individuals (Chesson, 2000). Nevertheless, the interaction coefficient assumes constant per capita density-dependent mechanisms. Thus, from a traditional competition perspective, an individual’s population size can only constantly decrease in the presence of neighbors. Alternatively, for a facilitative interaction, the net effect is restricted to increased facilitation with increased neighborhood density. Following such reasoning does not allow for the variation of species interaction, as would occur when species’ relationships with one another change across density, e.g. maybe facilitating at low density via shading and microhabitat modifications but competing at high density for space and limiting resources (McIntire and Fajardo, 2014).

#### 2.1.2 Alternative functional forms of species interactions

To include varying species interactions in the above model, we explore three alternative functional forms for estimating pairwise plant interactions, each allowing for variation in responses to neighbor identity and density. The functional forms we consider are linear, exponential, and sigmoidal relationships between the effect of per capita interaction and neighbors’ local density (fig.1b). The three functions represent a subset of possible mathematical function forms, but were selected because together they encompass a wide variety of different shapes depending on parameterizations and represent three additional levels of increasing complexity (Holland et al., 2002; Aguirrebengoa et al., 2023; Bruninga-Socolar et al., 2023).

The three functions we explore in this study introduce new parameters compared to the traditional constant net interaction outcome: the optimal neighborhood density (e.g., *N*_*o,i* *j*_), the density-dependent effect on interaction strength (e.g., α^_*i* *j*_) and the stretching parameter (e.g., α^^^_*ij*_). These new parameters allow variation in the effects of one individual of a species *j* on the intrinsic performance of a focal species, *i*, depending on the density of the *j* species. The optimal neighborhood density is the density of an interacting species for which its realized effect on the performance of a focal species is optimal, that is, minimal (closest to zero) for competitive species and maximal for facilitative species. These parameters and how they are embedded in the three new functional forms can be visualized in terms of their resulting functional shape and their impact on the prediction of the individual’s performance of a species *i* with the Shiny App available at Buchel9844/Density-dependentFunctions/shiny (release v1-, zenodo DOI:).

**Linear form** Under this functional form, the overall interaction effect is linearly dependent on neighbors-density (fig.1.b. and c. yellow function):

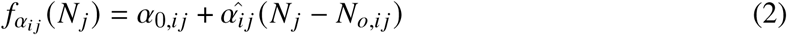

with α^_*ij*_ the density-dependent effect of *j* on *i* (α^_*ij*_ ∈ [−1, 0]), *N*_*o,ij*_ the optimal neighbors’ density of *j* when species *i* reaches maximal per capita performance. Here, the constant density-independent effect (α_0,*ij*_) defines the effect when the neighbors’ density is optimal, that is *N* _*j*_ = *N*_*o,ij*_. The density-dependent effect defines the slope, that is, the rate of change in the per capita effect of species *j* on species *i*, which can change drastically (high value of |α^_*ij*_ |) or stay stable (|α^_*ij*_ | ∼ 0) along the neighbor’s density gradient. The linear form follows a steady increase in the net interaction effect on performance, capturing the impact of resource modification from a neighbor density as a constant, unlimited effect. Under this scenario, species interactions change, say from facilitative to competitive, proportional to neighbor density and without threshold effects.

**Exponential form** The second functional form we consider is a negative exponential following neighbors’ density (fig.1.b. and c. blue function):

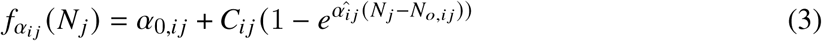

with α^^^_*ij*_ a constant responsible for vertical stretching (α^^^_*ij*_ ∈ [−1, 0]). The negative exponential functional form allows for rapid change at low density while approaching a lower asymptote, preventing excessively high competitive effects at high density. The density-dependent effect (e.g., α^_*ij*_) describes the shape of the exponential with a less or more pronounced asymptote. The stretching parameter (e.g., α^^^_*ij*_) describes the values of the asymptote, that is, the boundaries of the magnitude of the interaction effect. The net interaction outcome of *j* on *i* saturates at the asymptote when the neighborhood reaches the optimal density (i.e., *N* _*j*_ = *N*_*o,ij*_). The exponential form follows a fast increase in the net interaction effect on performance at low neighbor density until reaching an asymptote, capturing the potentially important effect of microhabitat modification when a neighbor first invades. For instance, an invader can drastically change the environment at the invasion stage and can benefit native species until reaching a higher density, where the invader then exploits native resources, leading to competition (Zarnetske et al., 2013). This functional form captures that the facilitative effect (or competitive release effect) on performance can be drastic at low-neighbors density, given the lack of upper bound.

**Sigmoid form** The final interaction form we consider is a sigmoid shape following neighbors’ density (fig.1.b. and c. green function):

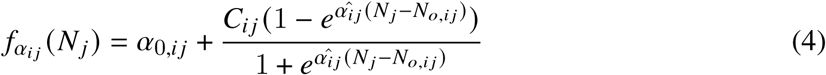

The sigmoid functional form introduces the same parameters as the previous, exponential functional form. A sigmoid form provides lower and upper boundaries to secure realistic interaction values at both low and high neighbor densities, preventing excessively high facilitative and competition effects, respectively. The net interaction outcome of *j* on *i* can switch relatively fast when the neighbor reaches the optimal density (i.e., *N* _*j*_ = *N*_*o,ij*_). For instance, the effect of shading from a grass species on a forb species might be fairly consistent until it reaches a density where it then begins crowding out the forb, yielding a narrow neighborhood density where a switch from facilitation to competition emerges. After this threshold, competition for resources or space may not intensify strongly with further increases in neighbor density.

### 2.2 Simulated communities From species interactions to population trajectory and community dynamics

We simulated density over time (i.e., time-series) for two species communities to compare the traditional function (i.e., a constant value of interaction) against the three alternatives. Based on the time-series, we assessed which functional forms produced population(s) that are persisting (i.e., positive growth rates at low density) and/or temporally stable (i.e., ratio of mean density over variance). Finally, we examined the impact of varying interactions on predicting oscillatory and/or synchrony dynamics in the coexisting simulated communities.

#### 2.2.1 General framework

We simulated densities over time for two species communities based on a generalizable individual performance model. The individual performance model evaluates the performance of species *i* through time t as the annual growth rate 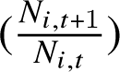. We defined it following the classic Ricker model with our four alternative species interaction forms incorporated as follows (Ricker, 1954):

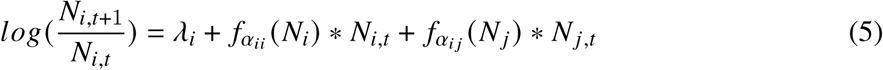

with γ_*i*_ the intrinsic growth rate and *f*_α*ij*_ (*N* _*j*_) the functional form of the net interaction outcome of *j* on *i*. The net interaction outcome is traditionally noted α_0,*ij*_ (see eq.1), yet it is now expressed as a function of neighbors density (noted hereafter *f*_α*ij*_ (*N* _*j*_)). Our new functional forms can be embedded in the individual performance models by respectively defining the *f*_α*ij*_ (*N* _*j*_).

Using the above individual performance model (eq.5), we simulated two-species communities for 200 time steps, using starting densities of *N*_*i*_ = *N* _*j*_ = 5. We simulated 4, 500 communities— 1500 per each of our four functional forms. To do so, we randomly drew 1500 sets of parameter values from the following distributions: α_0,*ij*_ ∼ *N* (0, 0.2), α^_*ij*_ ∼ *U*(−1, 0), *N*_0_ ∼ *U*(0, 10), α^^^_*ij*_ ∼ *U*(−1, 0) and α_0,*ii*_ ∼ −*abs*(*N* (0, 0.2)), where *N* denotes a normal distribution, *U* a uniform distribution, and *abs* the absolute value. We then used each set of parameter combinations to simulate dynamics according to the four function forms of net interaction outcome. That is, each parameter draw was used as input for functional forms 1, 2, 3, and 4 (appendix, section A.1.1).

Additionally, we evaluated the importance of facilitation on population trajectory by categorizing each set of parameters. We separated the 1500 sets of parameters for each functional form into three categories based on the predominant inter-specific interactive effect (i.e., the sign of the density-independent effect): (i) mainly inter-facilitation (α_0,*ij*_ > 0 and α_0, *ji*_ > 0, example on fig. 2.a), (ii) mainy inter-competitive (α_0,*ij*_ < 0 and α_0, *ji*_ < 0, example on fig.2.b) or (iii) one species is mainly facilitative and the other one mainly competitive (α_0,*ij*_ < 0 and α_0, *ji*_ > 0, or vice versa). Here, we consider the density-independent intraspecific effect to be only negative (i.e., α_0,*ii*_ ∈ [−1, 0]) as facilitation emerges primarily from inter-specific interactions (Adler et al., 2018). Yet, note that the net intraspecific interactions can still be positive at low density with the alternative functional forms, depending on the parametrization.

**Figure 2:**
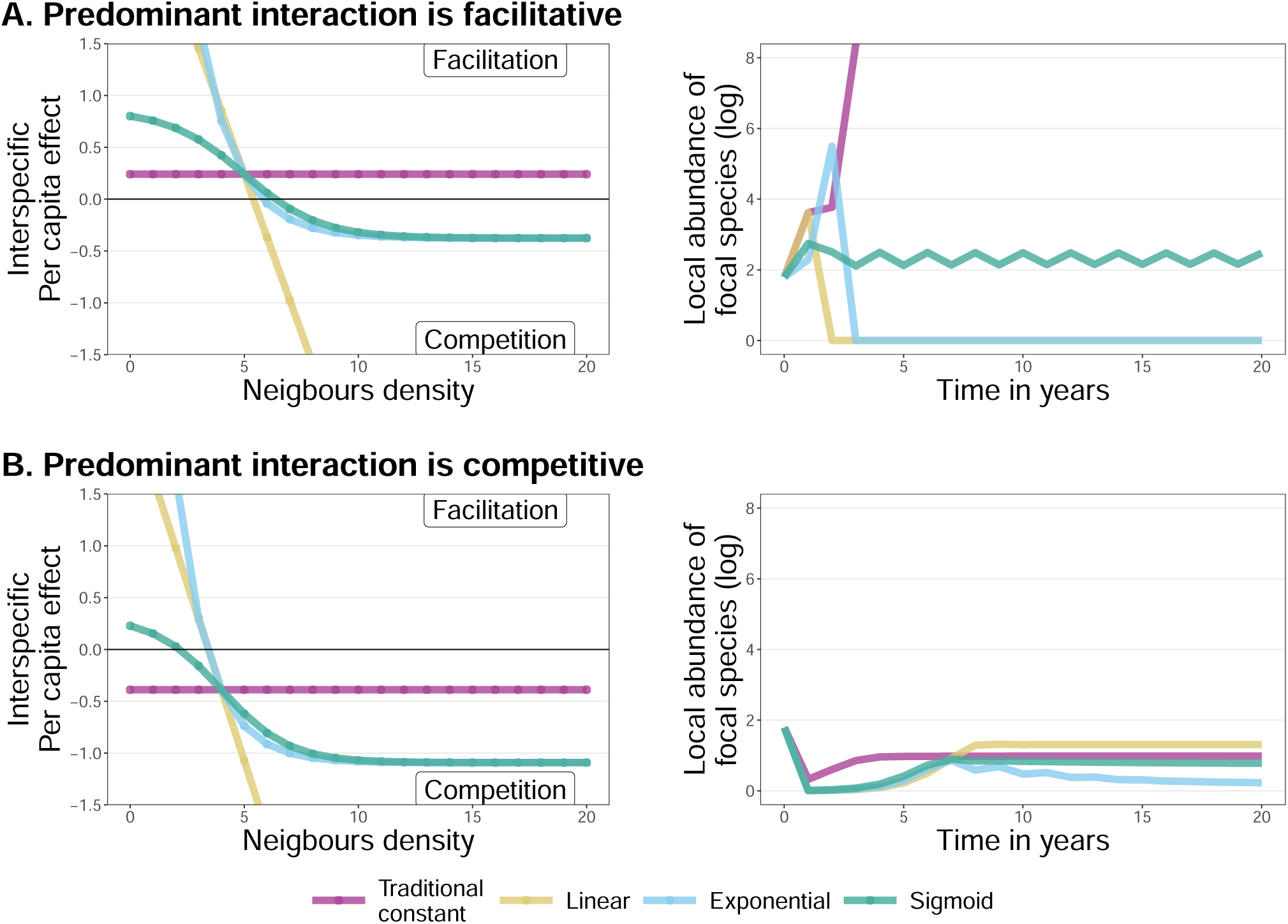
Example of the four functional forms (on the left) and their respective simulated densities over time (on the right) under two scenarios of predominant interactions (by rows). **Row a.** Shows an example of one set of parameters with predominant inter-facilitation, with the four functional forms on the left and the corresponding simulated density over time on the right. **Row b.** Shows an example of one set of parameters with the predominant inter-competition. The community dynamics are described by a Ricker model, eq.7. We visualize 30 years out of the 200 simulated.

**Figure 3:**
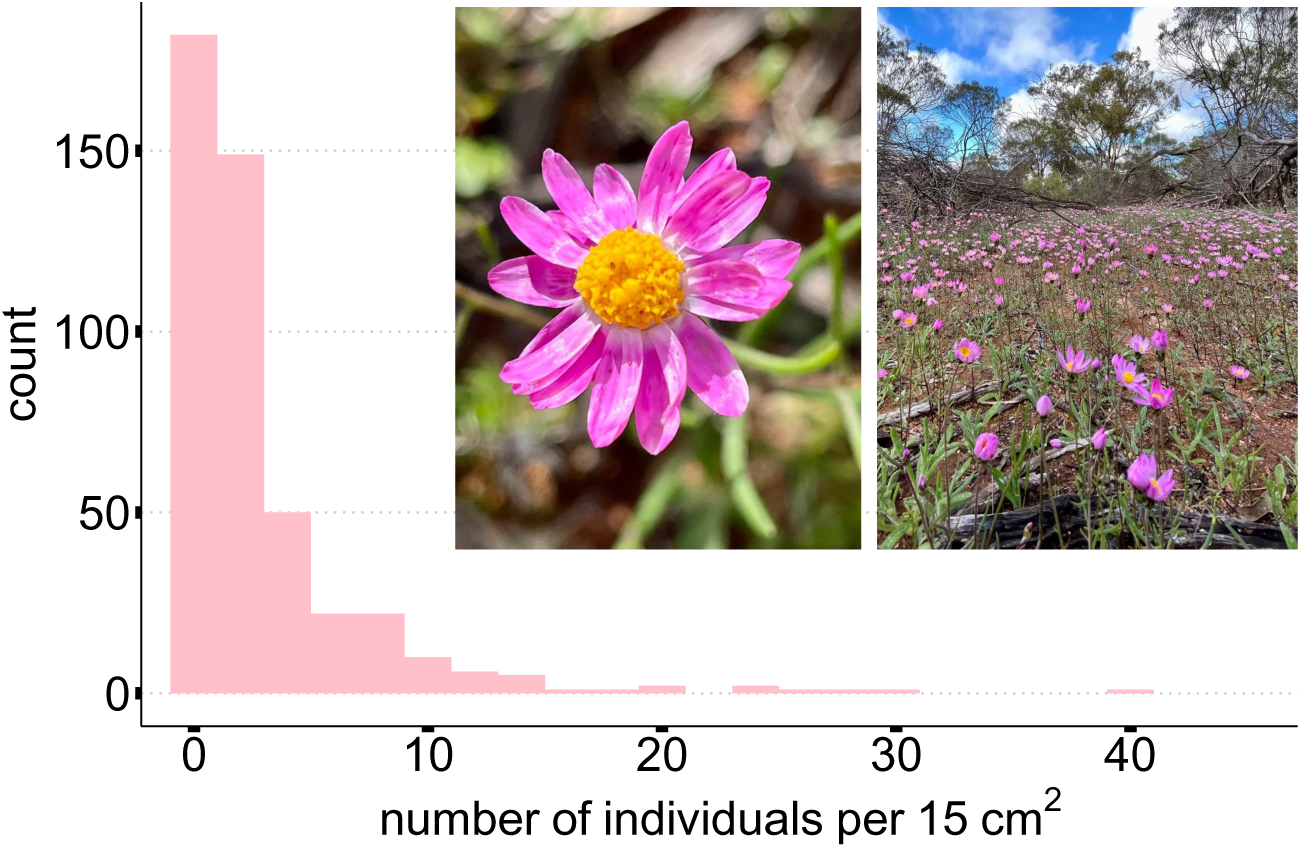
(a) Distribution of the number of *Lawrencella rosea* (LARO) per 15 *cm*^2^ observed across years. (left) The capitula of LARO; (right) A community of LARO in a York gum-Jam woodland, Perenjori, Australia. Credit to Wingman Siu.

#### 2.2.2 Evaluation of population trajectory

Across all simulated communities, we used population growth rates and densities to compare population persistence and temporal stability across the four functional forms. We evaluated the population persistence based on the growth rate when low (GRWL, rather than the invasion growth rate; see appendix, section A.3). To do so, we examined the growth rates at the lowest density of either species, excluding the first 99-time steps as a burn-in to minimize any potential influence from the initial values. In each community, we characterized the populations’ density over time as either (i) both species persist (GRWL for both species > 0), (ii) only one species persists (GRWL for at least one species > 0), (iii) neither persists (GRWL for both species < 0), or (iv) at least one species shows a runaway trajectory (GRWL for at least one species > 100). In addition to evaluating the growth rate when low, we also categorized the population trends by examining the median, maximum, and minimum densities reached during the time series, again excluding the first 99 time-step burn-in. (see appendix, tab.S2).

Additionally, we evaluated the temporal stability of populations using the ratio of mean density over variance (inverse coefficient of variation, Doak et al. 1998). A population with a ratio higher than one exhibits robustness over a long time frame, which we call temporal stability. High values of the ratio refer to high temporal stability as the mean density is much bigger than the variance (i.e., 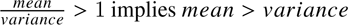. Inversely, a population with a small ratio (i.e. 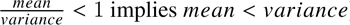) captures unstable temporal dynamics with a variance bigger than the mean density. A community of two species can be composed of (i) no stable populations, (ii) one stable population, or (iii) two stable populations.

#### 2.2.3 Prediction of community dynamics

Finally, we evaluated how fluctuations in the two populations’ densities relate to one another over time across the four functional forms of species interactions. That is, we examined if different functional forms produce different internally driven temporal dynamics. We differentiate between oscillatory dynamics, which defines if the mean density of a population oscillates over time, and synchrony, which quantifies if these oscillations are correlated between the two populations. Synchrony can occur over short-term and/or long-term time scales (e.g., high-frequency, annual synchrony versus low-frequency oscillations over multiple generations) (Zhao et al., 2020; Shoemaker et al., 2022). Thus, we categorized community dynamics for each simulated community that was predicted to have two persisting species, based on three criteria representing natural dynamics observed in annual plants: (i) oscillation (function auto.arima from the package *forecast version 8.21*, Hyndman et al. 2023; Turchin and Taylor 1992; Berryman and Lima 2007; Plaza et al. 2012, (ii) short-term and (iii) long-term synchrony (function tsvreq_classic from the package *timescale-specific variance ratio version 1.0.2*, Zhao et al. 2024; Shoemaker et al. 2022) (see appendix, tab.S5 for condition). The presence of one process is not mutually exclusive from the presence of another. Communities can also exhibit no specific dynamics if the populations do not fluctuate according to either oscillatory or synchronous patterns.

### 2.3 Natural communities | Species interactions of *Lawrencella rosea*

We used a natural community of annual plants to show how we can improve our explanation and prediction of density over time for one population (*Lawrencella rosea*, which we shorten to LARO) based on its interactions with multiple neighbors. We used a competitive experiment to assess the effect of the main taxonomic families on LARO, fitting fecundity models with each functional form. Based on the estimated species interaction parameters from each model, we compared the predicted density of LARO over time with its past observed density.

*Lawrencella rosea*, (LARO) is a native annual Asteraceae abundant in woodlands around Perenjori, WA, Australia. Native remnants in the Perenjori region are dominated by the York gum-Jam woodlands (YGJW), which comprised *Eucalyptus loxophleba* Benth. (York gum) and *Acacia acuminata* Benth. (Jam) overstory, with a diverse understory of annual forms and grasses and perennial geophytes, tussock grasses and shrubs (Raymundo et al., 2021). The region has a semi-arid Mediterranean climate with hot, dry summers (25-36^◦^C) and mild, wet winters (5–17^◦^C) and a mean annual precipitation of 309 mm (90-year average) (*Bureau of Meteorology* 2023). This rainfall occurs primarily during the winter and spring months (June to October), when annual wildflowers germinate and grow. The summers are hot and dry with prolonged periods of low-to-no rainfall (November to March) (*Bureau of Meteorology* 2023). Previous studies have demonstrated that this system involves substantial numbers of positive species interactions (Bimler et al., 2018; Wainwright et al., 2019; Bowler et al., 2022).

#### 2.3.1 Quantifying plant-plant interactions

To quantify plant-plant interactions between LARO and its local neighbors, we measure the per germinant viable seed production (i.e., performance, measured as fecundity) as a function of both the number and identity of local neighbors (inter- and intra-specific) within a diameter of 15 cm. This radius is a standard distance used in previous studies to measure competitive interactions among annual plant species (Levine and HilleRisLambers, 2009; Mayfield and Stouffer, 2017), and has been validated to capture the outcome of competition interactions at larger scales (1*m*^2^) under locally homogeneous environmental conditions (Godoy et al., 2014). Seed production of focal LARO individuals and neighborhood composition were measured across 118 locations during one growing season. For model fitting, we grouped LARO’s neighbors into categories: conspecific individuals (LARO), individuals of abundant families (Araliaceae, Asteraceae, Crassulacea, Goodeniaceae, Montiaceae, Poaceae), and individuals of rare families (families with densities smaller than 10% of total densities). These groups differentiate neighbors with the potential for differing interaction strategies (Martyn et al., 2020). For instance, LARO likely interacts differently with con-specific individuals (i.e., another LARO) vs hetero-specific neighbors. Similarly, flowering neighbors (e.g., Asteraceae) compete for floral visitors, while non-flowering neighbors (e.g., Poaceae) interact for other resources.

We build on our simulation model methods to estimate the effect of each neighbor group on LARO’s fecundity, fitting the Ricker individual performance model (Ricker, 1954). We estimate the model parameters describing the per capita fecundity:

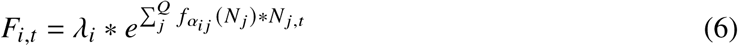

with *N* _*j*,t_ the number of observed stems of family *j* at time t, λ_*i*_ the fecundity of the focal species when growing alone. The per capita effect of each family was estimated by *f*_α*ij*_ (*N* _*j*_), which is defined by one of the functional forms: the traditional constant value (eq.1), the linear (eq.2), the exponential (eq.3) or the sigmoid (eq.4). The effect of each family is summed over the set of all neighbor groups *Q*. The intraspecific interaction (i.e., *f*_α*ii*_ (*N*_*i*_)) was not restrained to competitive interactions.

To fit eq.6 to the empirical data, we used a Bayesian approach in Stan, with *log*(*F*_*i*,t_) described by a negative binomial function. We fit each model using four chains with 3000 iterations and 1000 warmups; all models exhibited good effective sample sizes and convergence (The Stan Development Team, 2014). All code is available on https://github.com/Buchel9844/ Density-dependentFunctions and details are in the appendix A.5).

#### 2.3.2 Simulating time-series

After fitting the per capita fecundity (eq.6) for each functional form to the empirical data, we projected LARO’s density over the next 20 generations (i.e. time steps) using a classic annual plant model (Cohen, 1966), which describes the total number of seeds in the seed bank as:

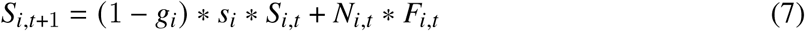

with *g*_*i*_ the germination rate, *s*_*i*_ seed survival rate of dormant seed (here *g*_*i*_ = 0.2 and *s*_*i*_ = 0.4, based on previous unpublished green house experiment), *N*_*i*,t_ the number of stem of species *i* at time t and *F*_*i*,t_ the viable seeds produced per germinated individual of focal species *i* at time t as determined from our fecundity model (eqn 6).

For our simulation of LARO’s densities through time, we based the future composition of LARO’s neighbors according to past field observations. The Mayfield lab recorded the local density of LARO and other locally present species in multiple locations in the Perenjori region from 2010 to 2023 (see appendix, Fig.S12, collected by Dwyer et al. (2015), Wainwright et al. (2016), Raymundo et al. (2021), James (2023), and Sevenello et al. (2024); Taylor et al. in prep). We summarized observations of neighborhood densities within each year by aggregating neighbors according to the family groupings mentioned above. Based on each family’s densities between 2010 and 2023, we extrapolated 100 potential time series scenarios from 2024 to 2043. First, we computed the residuals correlation matrix between each family’s density across the 13 years by fitting a multivariate Poisson to our observed time-series (PNL, *PLNmodels* package Chiquet et al., 2021; appendix, Fig.S15). The multivariate Poisson accounts for correlation patterns in density and allows us to maintain these patterns in the simulated time series (appendix, Fig.S13,S14). After we fitted the multivariate Poisson, we predicted the density of the neighbor families for the next 20 years conditional on the observed density of the Poaceae family (randomly chosen out of all the families), which was itself drawn from a univariate poison distribution based on its overall density (predict_cond, *PLNmodels* package Chiquet et al., 2021 and fitdistr, *MASS* package Venables and Ripley, 2002). We repeated this sampling to generate 100 potential time-series of neighbor densities. For each time-series, we estimated the temporal dynamics of LARO based on the densities of its neighbors and Eqs.6 & 7.

## 3. Results

### 3.1 Plant-plant interactions can vary according to neighborhood density and species identity

Varying species interactions according to the neighborhood density increases the number of parameters to estimate per pairwise interaction by three for the linear functional form and four for the exponential and sigmoid case. This increase in model complexity does not come at the cost of model convergence (tab.S8), suggesting that including variation in interaction strengths still allows for theory-data integration despite the larger number of parameters to estimate (see box.5 for mathematical assumptions and parameter examination). For the most complex functional forms, we also find good parameter recovery in a computational experiment (appendix, section A.6) and good parameter convergence when fitting empirical data, suggesting that they can be considered plausible forms of species interactions. Additionally, the models of each functional form have similar explanatory power (see appendix, leave-one-out outcome tab.S8, and post-predictive fits Fig.S11).

Regardless of the functional form evaluated, the neighborhood groupings that interact with *Lawrencella rosea*, LARO in the natural community follow a consistent order of interaction strength. At one extreme, individuals in the Goodeniaceae family compete in all instances, while at the other extreme, individuals in the Crassulaceae have the most positive interactions (fig.4). All families except Crassulaceae compete with LARO when modelled using the traditional constant function. Yet, they all show low-density facilitation with the other functional forms, except individuals of conspecifics (LARO grouping) and the Goodeniaceae family for the exponential and sigmoid forms. The range of values adopted by the interactions using the traditional form is extremely weak (*f*_α*ij*_ (*N* _*j*_) < 0.1), the linear forms reach extremely strong values (*f*_α*ij*_ (*N* _*j*_) > ∓1), while the sigmoid and exponential have a similar range (−0.2 < *f*_α*ij*_ (*N* _*j*_) < 0.2). Despite these differences between the range of values, the families’ net effect on LARO (when interaction effects are multiplied by each family’s observed abundance, appendix, Fig.S10) is comparable between functional forms. LARO experiences the most competitive effect from the Asteraceae family, which is the most abundant family of the system. Yet, according to the sigmoid and exponential functions, individuals from the Crassulaceae, the Montiaceae and Poaceae families tend to positively impact LARO as they are most often present in low abundance in LARO’s neighborhood.

**Figure 4:**
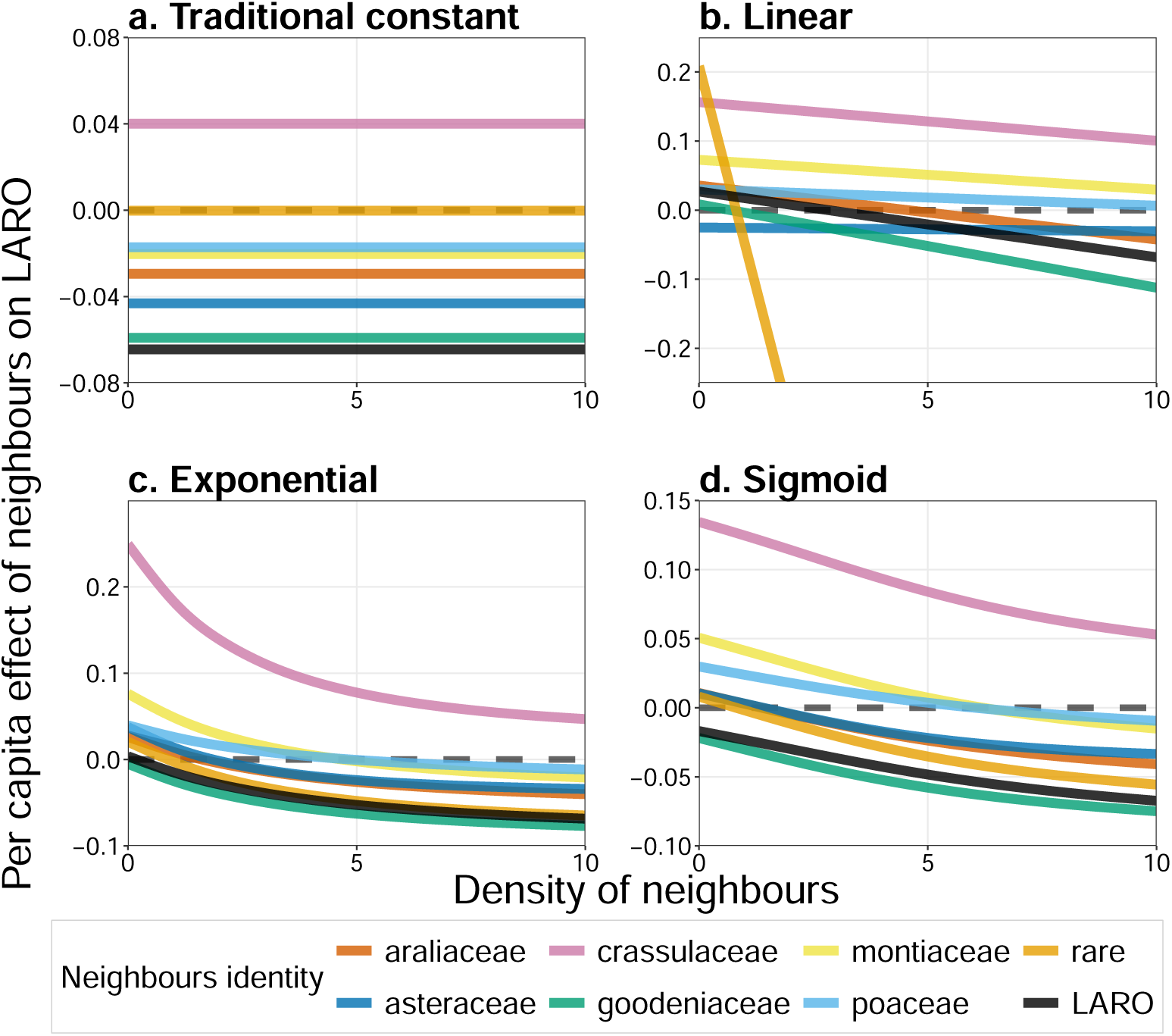
Empirical data – The effect of the main families present in the natural community on *Lawrencella rosea*, LARO, according to each functional form. The per capita effect (y-axis) of individuals of a family on LARO can change according to the family’s density (x-axis) when using the linear, exponential or sigmoid functional forms. Note, scale of y-axis varies between panels.

### 3.2 The sigmoid functional form shows the most realistic population trajectories in our simulated and empirical datasets

More specifically, in our simulations, out of the 1500 two-species communities simulated for each functional form, we find different percentages of coexisting communities (positive GRWL): 378 communities for the traditional form (25.3%), 222 communities for the linear form (14.9%), 331 communities for the exponential form (22.2%), and 857 communities for the sigmoid form (57.4%) (see first row, Fig.5, and appendix, tab.S3). Therefore, coexistence is the most common result of the sigmoid relationship, and the probability of coexistence across the parameter ranges we considered is doubled with the sigmoid form compared to all other functional forms. While predominant facilitation yields dominant runaway dynamics (i.e., unbounded increase in abundance) or extinction with all of the other forms, it primarily allows for coexistence with the sigmoid form (64.8%, fig.5, second row). Indeed, the traditional functional form, which has a constant value for interaction effects, is the most likely to predict runaway population growth (26.2%). When interspecific facilitation is more common than competition (α_0,*ij*_ > 0 and α_0, *ji*_ > 0; i.e. predominant facilitative community), the traditional functional form shows mainly runaway dynamics (76.4% of all facilitating communities), unlike in all other functional forms, for which this is a low probability outcome. In contrast, the linear and exponential functional forms resulted in more extinctions (73.8% and 49.7% respectively, tab.S3), regardless of the predominant sign of interspecific interactions.

**Figure 5:**
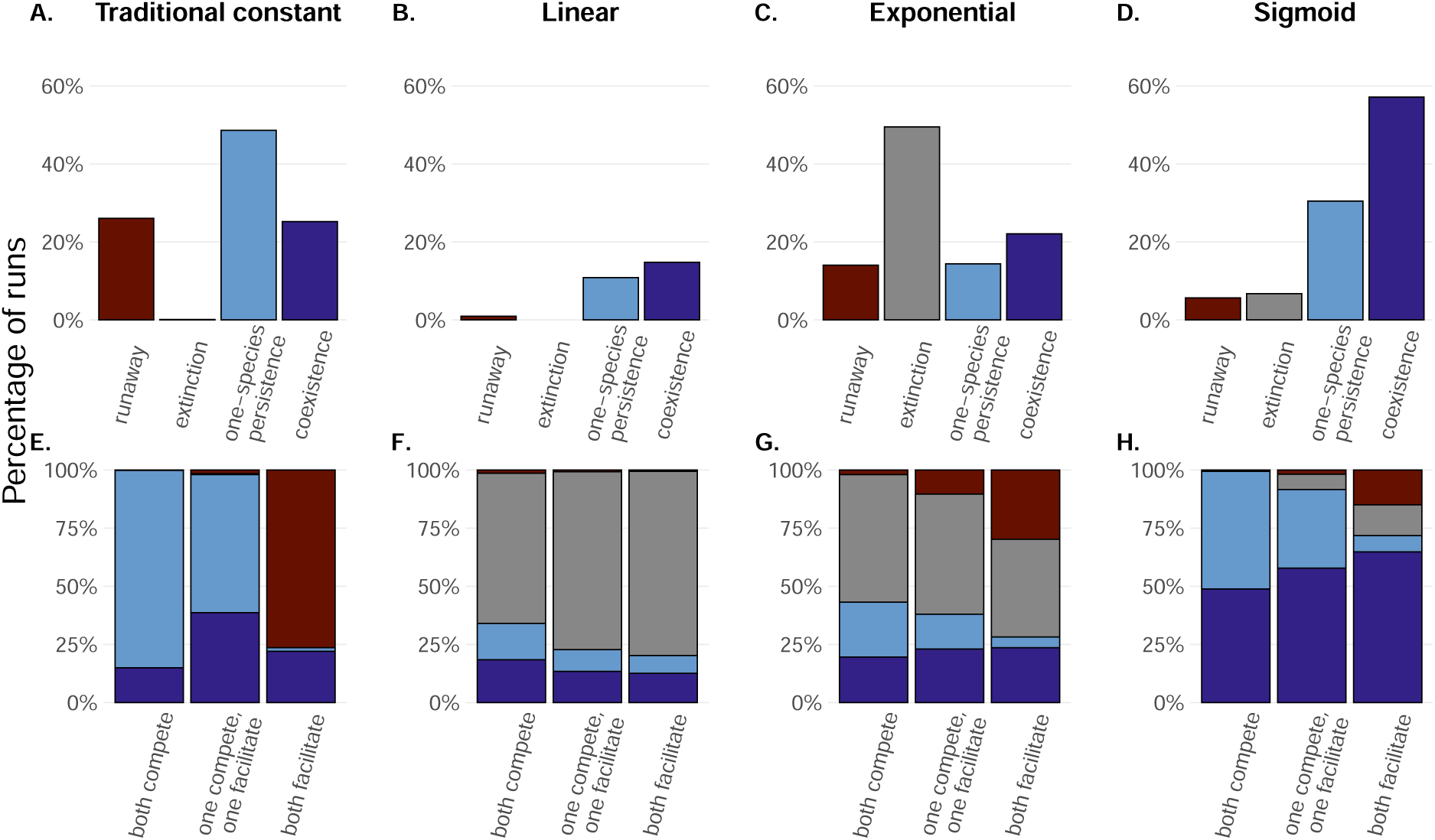
Simulated data – Population outcomes differ based on the functional form and the predominant sign of the interspecific interactive effect. The **first row** shows the proportion of each community composition predicted according to each functional form. The **second row** shows community composition from the top row, aggregated according to the predominant sign of interspecific interactions: either mainly inter-facilitation between both species, mainly inter-competition between both species or one species mainly facilitative and the other mainly competitive).

In the empirical system, both non-linear functional forms, exponential and sigmoid produce more realistic population abundances of *Lawrencella rosea* (LARO) over time, with model results best matching empirical patterns. From 2010 to 2023, the observed abundance of LARO had a mean of 1.84 and standard deviation of ±3.68. The traditional constant (scalar) functional form predicts a higher LARO density, with peaks reaching about 66 individuals and an average of 10.40 ± 8.12. (Appendix, tab.S7). This range is overestimated compared with the observed mean population size (fig.6.b). According to the linear function, the local population density dropped to an average of 0.79 ± 2.50 and a median of 0.002, predicting the extinction of LARO, which we did not observed in our system. The exponential and sigmoid best predict the local population density with a mean equal to 5.32 ± 5.70 and 6.67 ± 6.48, respectively, and density’s distributions similar to LARO’s observed density over time (fig.6).

**Figure 6:**
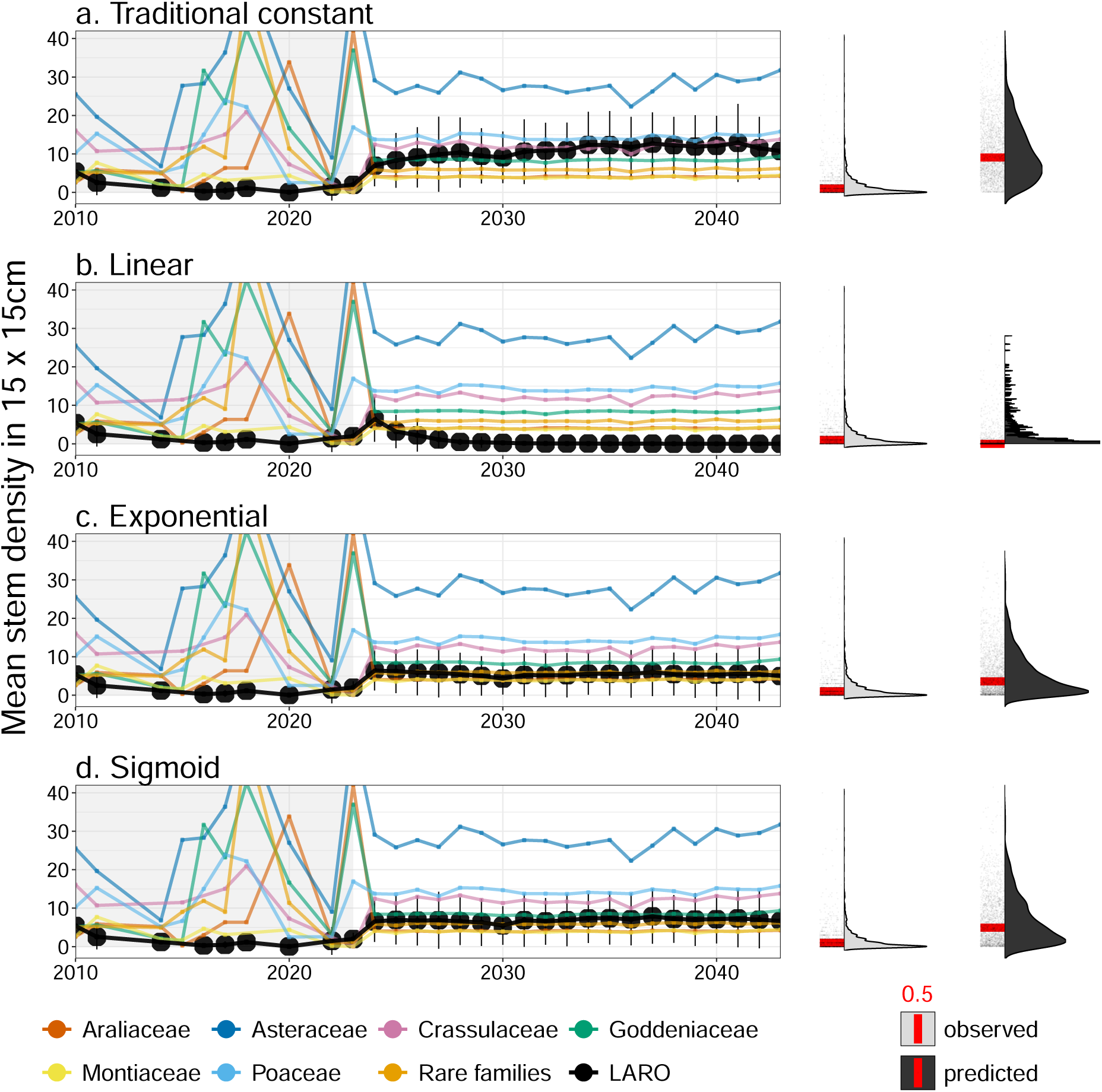
Empirical data – Including varying interactions with a bounded functional form, such as an exponential or sigmoid, leads to more realistic prediction of local abundances for LARO. As the effect of each family differs based on the functional forms, the prediction of LARO’s local abundances over the next 20 years also varies based on how we estimate species interactions. For each time step, each LARO’s dot represents the mean and the standard deviation over the 100 predictions (left column). The neighboring families only show the mean for each time step. The time series distinguishes the observed data (in the grey area) and the predicted data >2023 (in the white area). The density distribution of the observed (grey) and predicted (black) LARO can be compared on the right side, with the median in red (previously observed density: 0.5; Traditional constant: 8.56; Linear: 0.00; Exponential: 3.372 and Sigmoid: 4.64) and the raw points below the distributions.

### 3.3 The sigmoid functional form yields a diversity of community dynamics, including oscillatory and synchronous dynamics

In the simulations, the sigmoid and traditional constant functional forms have a similar percentage (∼ 21%) of coexisting communities predicted to be stable (fig.7, and appendix, tab.S4). For communities where both species did not show temporal stability (i.e., the ratio of mean over variance < 1), the sigmoid functional form shows a wider diversity of community dynamics, such as short or/and long oscillation (10.3%) and synchrony (7.4%), while coexisting. Across all communities, the sigmoid form produces 25.5% of communities with at least one form of internal dynamics. In contrast, the percent of communities showing oscillation and/or synchrony is very low across all other functional forms: 2.8% for the traditional constant functional form, 3.5% for the linear functional form and 6.9% for the exponential functional form (appendix, tab.S4), regardless of coexistence and stability regime. The exponential and the sigmoid forms show similar proportions of communities with temporal stability, yet the exponential form had fewer coexisting communities overall. Thus, we find strong evidence that varying the strength of species interactions according to the sigmoid functional forms, allows for obtaining synchronous temporal dynamics driven by internal oscillations while maintaining species persistence and community coexistence.

**Figure 7:**
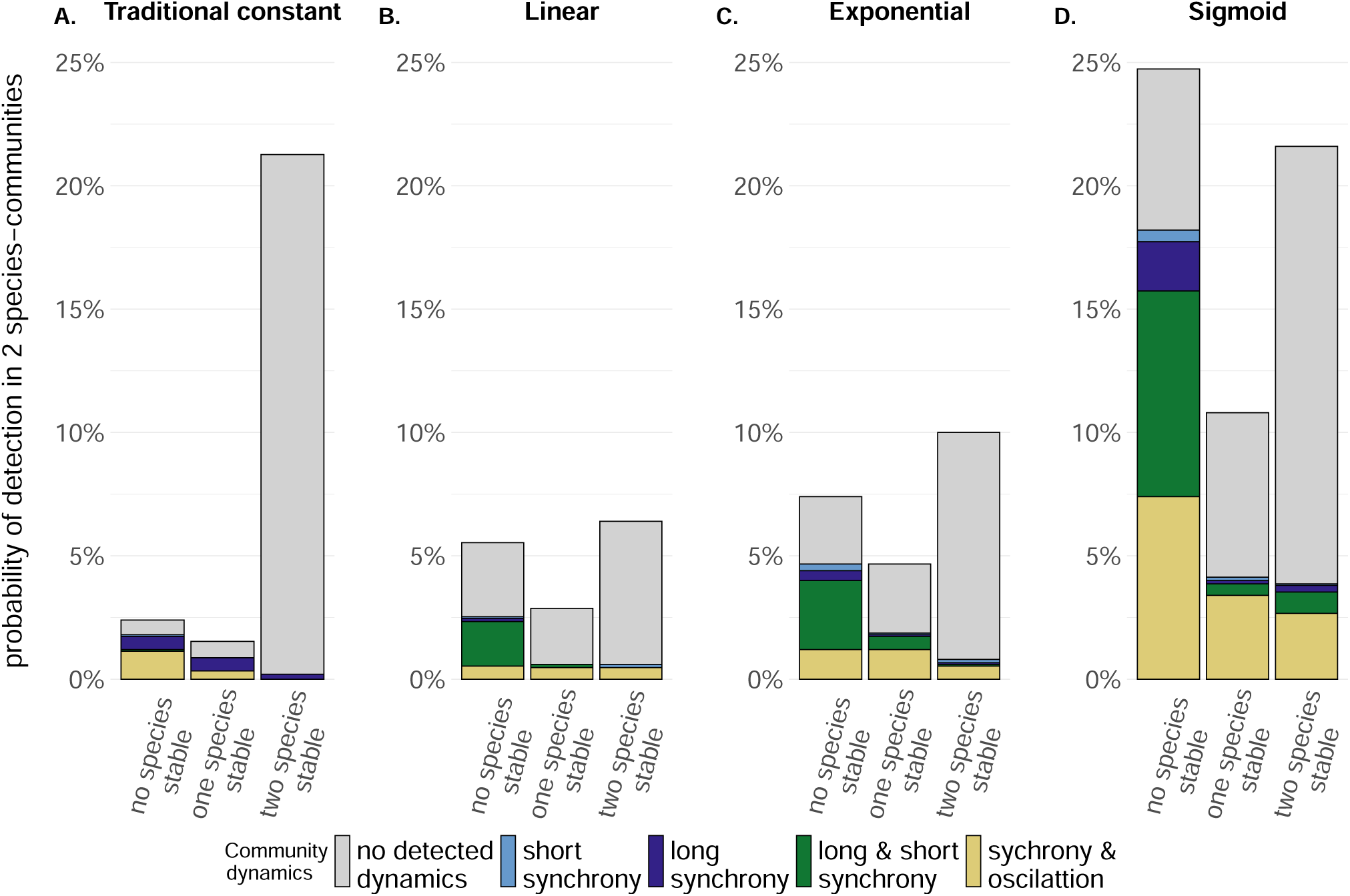
Simulated data – Different functional forms can lead to diverse community dynamics; the sigmoid form has the most variable outcome. The bars don’t add up to 100% but add up to the probability from Figure 4 of having a 2-species coexisting community (appendix, tab.S3). We categorised each coexisting community according to whether both species were temporarily stable, only one out of the two, or either. Colours show predictions of different internal dynamics: long and/or short synchrony, and oscillation.

## 4. Discussion

Our simulations and investigations of an empirical community show that including non-linear effects of neighbor density on interaction strengths improves predictions of population persistence, stability, and community dynamics. Analyses of both simulated and empirical studies revealed that the bounded sigmoid functional form was the best function for balancing natural complexity and mathematical feasibility. Using the sigmoid function increased the persistence and coexistence of species in communities, yielding internal fluctuations in abundances, and bounding species’ abundances to realistic values. This functional form also allowed for facilitation while preventing ‘run-away’ population growth. As such, we argue that the oversimplifications of typical interaction models to a single, static interaction strength should be reconsidered for models that use dynamic functional forms.

### 4.1 Varying species interactions outcome

We show, for the first time to our knowledge, that allowing the net species interactions to vary from facilitation to competition across neighborhood densities leads to more realistic population trajectories than when constraining it to a static, usually competitive, interaction strength regardless of density. Estimates of species interaction strengths encompass multiple ecological processes, including microhabitat modification, competition for multiple limiting resources such as water, soil nitrogen, and soil phosphorous, and attraction of pollinators. As such, it stands to reason that the net interaction might differ across neighbor densities. Incorporating this context dependency is an improvement to past modeling efforts that typically deal with facilitation by restricting it artificially or by averaging to ensure that overall outcomes remain competitive (fig.7 and fig.6) (Hart, 2023; Bertness and Callaway, 1994; Bertness and Hacker, 1994; Doak et al., 1998). Our simulations of constant facilitation across neighbor density resulted in runaway dynamics as expected. When we used a bounded sigmoid shape, however, we found more persisting species and increased pairwise coexistence without runaway facilitative dynamics.

In our analyses of *Lawrencella rosea*, the sigmoid and exponential functional forms predicted population abundances similar to the observed mean values while remaining above zero. Constant and linear functional forms, however, predicted patterns quite different from those observed in our field system. We hypothesize that the improved predictions resulting from the sigmoid and exponential functions reflect that these functional forms allow for more context-dependent processes to be captured. For this system, context-dependent processes likely include gradients in multiple key environmental factors including shading from the tree canopy and soil phosphorous (Bimler et al., 2018; Wainwright et al., 2019; Raymundo et al., 2021), demographic variation (Bowler et al., 2022), climate change (Dwyer et al., 2015) and annual variation in pollinator abundances (Loy et al., 2015; Sevenello et al., 2024). These context-dependent processes do not act in isolation but have a cumulative effect on plant performance (individual survival, fecundity, growth), which is observable as changes in plant density.

### 4.2 A more holistic understanding of species coexistence

Current theories of biodiversity maintenance are based on the phenomenological principle that species are more likely to coexist when intraspecific competition is greater than interspecific competition (Chesson, 2000; Saavedra et al., 2017; Barabás et al., 2018; Spaak et al., 2021). This principle aligns well with empirical observations from a wide diversity of ecological systems (Adler et al., 2018; Buche et al., 2022). Despite this conceptual alignment, efforts to predict coexistence in naturally complex systems often fail (e.g., Matias et al. 2018; Bimler et al. 2018). This mismatch between observation and prediction could be due to multiple missing sources of key information such as environmental variation (Bimler et al., 2018), or feedback dynamics across trophic levels (Lanuza et al., 2018; Buche et al., 2024). Another source of error could be the exclusion of variation in species interaction strengths in coexistence models. Our results show that when species interactions are allowed to vary with neighbor density, we predict more cases of pairwise species coexistence than we do with typical coexistence models. Other studies, such as by (Hatton et al., 2024), have used sublinear growth rate models to improve the fit of predicted and observed coexistence. While these models do a good job predicting biodiversity maintenance, the ecological mechanisms behind this maintenance remain unknown (Aguadé-Gorgorió et al., 2024; Letten, 2024). Our approach fits well within a slightly expanded version of the existing ecological theory that is typically used to understand coexistence in plants. Notably, our results are consistent with coexistence resulting from variation in how species respond to limiting resources (e.g. space, pollinators, water) through modifications in their interactions with other species. The only modification that our modeling approach requires is for coexistence to be possible when facilitation and competition are balanced across intra- and interspecific interactions and along density gradients. This balance for coexistence is supported by evidence from all of the functional forms evaluated in this study (box.5).

Our results require a deep consideration of two novel ideas. First, findings from both simulations and empirical observations provide strong evidence supporting the adoption of a sigmoid functional form for species interactions in studies of coexistence and diversity maintenance. The sigmoid function consistently and strongly reproduces a wider range of population dynamics than typical static models and better describes those dynamics in nature. Second, our results require some reconsideration of current coexistence theory. An easy step forward would be to start with coexistence framework such as modern coexistence theory (Chesson, 2000; Barabás et al., 2018) but expand the concepts of niche and fitness differences to include variation in interactions and all for more than two-species to interact at a time (Spaak et al., 2023). Another possibility would be to use the mutual invasibility framework of coexistence (Grainger et al., 2019; Hofbauer and Schreiber, 2022), along with non-traditional mechanisms of coexistence (Johnson et al., 2022; Levine et al., 2023; Zou et al., 2024; Godoy et al., 2024), or multiple alternative stable states in which species coexistence depends on the interplay between asymptotic and transient dynamics (Hofbauer and Schreiber, 2022; Spaak and Schreiber, 2023; Song1, 2024; Godoy et al., 2024).

### 4.3 Population and community outcomes

Oscillatory, synchronous, and otherwise temporally variable dynamics are ubiquitous across time series of abundances (Vasseur et al., 2014; Shoemaker et al., 2019; Barraquand et al., 2022; Firkowski et al., 2022). We often assume that these fluctuations are driven by environmental variation (Tredennick et al., 2017), but our results suggest that this is not always the case. This type of dynamic may also be driven internally by variations in species interaction strengths. For example, theory suggests that oscillatory dynamics can result from over-compensatory population growth (i.e. large density-independent growth rates that cause populations to overshoot their carrying capacities) (Turner et al., 2016; Grosholz et al., 2021). Our results suggest that these dynamics can also result from fluctuations in interaction strength, though this mechanism has been infrequently studied. As such, we suggest that future empirical work in highly controlled microcosms, could test for temporal oscillations and the relative sensitivity of dynamics to environmentally-driven effects, effects of density-independent growth rates yielding overcompensatory dynamics, versus variation in interaction strengths driving oscillations.

In our simulated two-species communities internal temporal dynamics depended on the functional form that we used (Fig. 7). The sigmoid functional form commonly produced both oscillatory and synchronous dynamics in temporally stable communities. Synchrony is often assumed and indeed is mathematically related, to stability, with more synchronous communities assumed to be less stable (Loreau and Mazancourt, 2013; Shoemaker et al., 2022). Yet we found that both oscillation and synchrony can co-occur in stable or partially stable communities if the interaction effects between individuals vary with one (i.e., exponential function) or two asymptotes (i.e., sigmoid function). These asymptotes can mediate systematic compensation for variation in population abundance; this then allows for temporal stability while oscillatory or synchronous dynamics are present (Loreau and Mazancourt, 2013). These asymptotes also allow a population subject to strong, direct, intraguild facilitation to be stable; a result that has been empirically observed (Sanmartí et al., 2022), yet never (to our knowledge) theoretically described (but see Kéfi et al. 2007; DeAngelis et al. 2012; Kéfi et al. 2016; Danet et al. 2021 for the effect of indirect facilitation on stability).

In non-temporally stable communities, we observed a high probability of detecting other dynamics, such as long-term and short-term synchrony, when communities are structured using a sigmoid form for interactions. The most coexistence occurred when we modeled using the sigmoid functional form. These coexisting communities were not stable but they were governed by oscillatory and synchronous dynamics. A study by Stouffer et al. (2018) previously showed how coexistence could be achieved in non-equilibrium circumstances by the presence of cyclic dynamics. Here, we provide evidence that other internal dynamics can be present and potentially responsible for the coexistence of non-stable populations.

### 4.4 Conclusion

Using simulations and empirical data, we showed that allowing the strength of species interactions to vary across densities reconciles empirical evidence of facilitative interactions with foundational theoretical models of coexistence. This presents an opportunity to reevaluate past conclusions about the prevalence of competition in natural communities (Yang et al., 2022), and provides a more efficient and effective tool for understanding and predicting the complex dynamics governing natural communities. Our findings strongly support the use of varying functional forms for estimating species interactions in studies aiming to understand biodiversity in multi-species communities. We suggest that future experiments explicitly consider the variation in the density of natural neighborhoods when attempting to predict coexistence in nature. We hope our findings will motivate theoretical studies of the role of facilitation and variation of species interactions in driving community diversity.

## 5. Box: Parameter sensitivity

The introduction of new parameters in alternative functional forms of species interactions allows the consideration of more natural complexity. It opens the question of the impact of these parameters on the community’s temporal stability. Using generalized linear models, we evaluated the relationship between the parameters within each functional form and temporal stability; fig.8).

**Figure 8:**
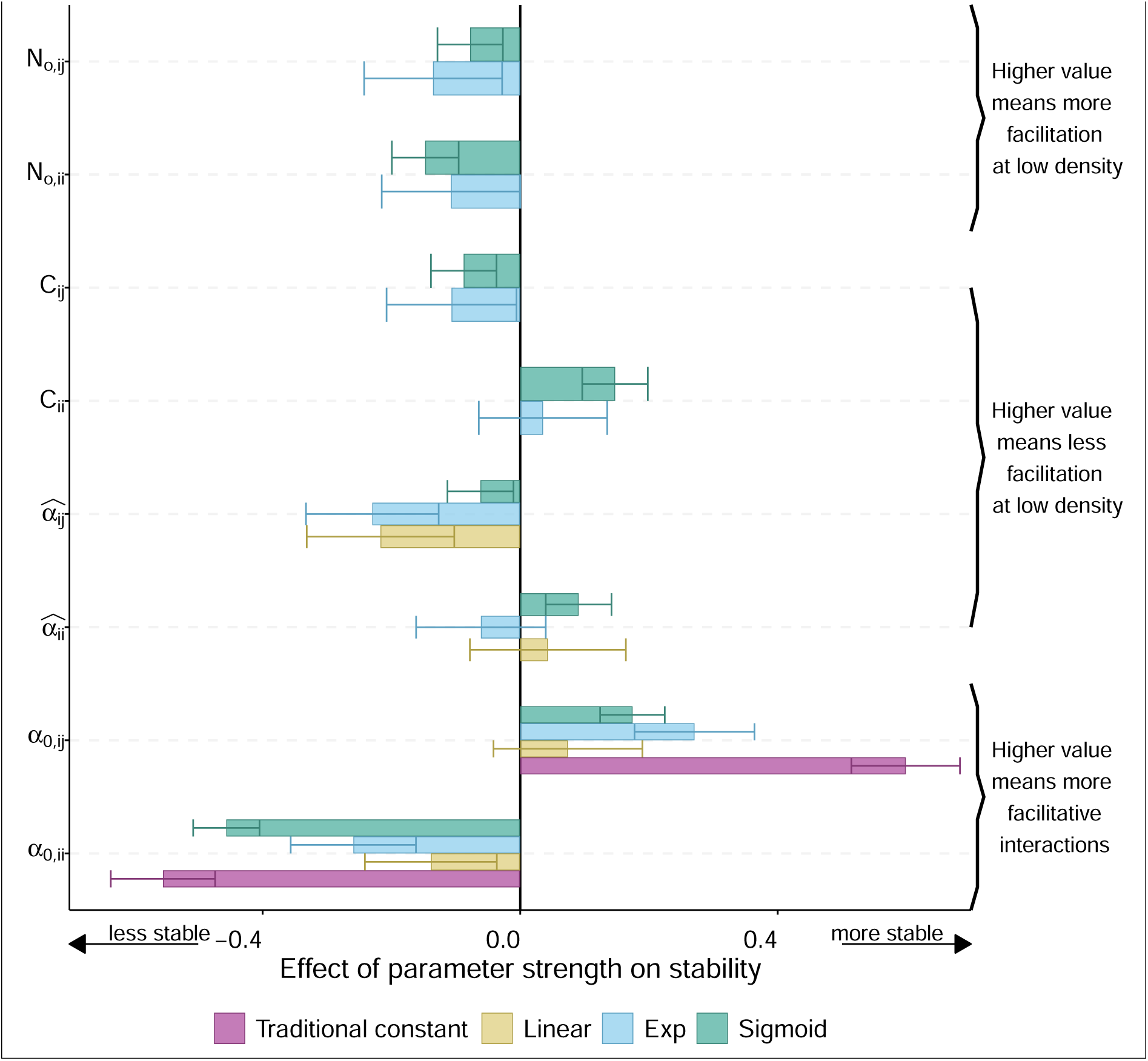
Parameters effect size on temporal stability with standard error. The expectation of high values of intra-specific and inter-specific parameters having a positive and a negative correlation with temporal stability respectively is respected for the density-independent effect (i.e., α_0_).

Following our expectations, we found that lower competitive intraspecific density-independent effect (small value; i.e., α_0,*ii*_, with range [−1, 1]) negatively impacts temporal stability, and this result holds across all functional forms. Indeed, if intraspecific interactions (i.e., α_0,*ii*_ > 0) do not compensate for the potential positive inter-specific effects, the chance of having stable populations decreases as populations can self-facilitate. By contrast, interspecific α_0,*ij*_ density-independent effects are proportional to temporal stability because positive interspecific effects favor a positive growth rate when low and, therefore, the maintenance of species.

When density-dependent effect (α^_*ij*_, with range [−1, 0]) is small, it increases the chance of seeing facilitation at low density. Thus, intraspecific density-dependent effects (α^_*ii*_) are positively proportional with temporal stability: stability is more probable if intraspecific effects have a lower chance of having facilitation at low density, which happens when the density-dependent effects are large (closer to zero than more negative); and inversely for the inter-specific density-dependent effects (α^_*ij*_). We see a similar behavior for the stretching parameter (α^^^, with range [−1, 0]) where intraspecific effects (α^^^_*ii*_) have a positively proportional effect on stability. Indeed, a position closer to zero promotes competition rather than facilitation at low density. Finally, high optimal density (*N*_*o*_, with range [0, 10]) translates to more facilitation at low density, negatively impacting temporal stability. Overall, we see that temporal stability at a local scale can be achieved if facilitation and competition are balanced across intra- and interspecific parameters. That is, a species’ chances to persist are based on how much it favors self-facilitation compared to how much it suffers from interspecific competition at high density.

## Acknowledgements

The authors thank Malyon Bimler for comments on an earlier version of the manuscript. We want to thank John Dwyer, Claire Wainwright, Maia Raymundo, Trace Martyn, Victoria Reynolds, Catherine Bowler, Aubrie James, Abigail Pastore, and Manuel Sevenello for the data collected in the Perenjori region. LB acknowledges financial support provided by the Holsworth Wildlife Research Endowment – Equity Trustees Charitable Foundation & the Ecological Society of Australia. OG acknowledges financial support provided by the Spanish Ministry of Economy and Competitiveness (MINECO) and by the European Social Fund through the TASTE project (PID2021-127607OB-I00). LGS was supported by NSF grants 233292 and 2019528. LMH was supported by NSF, Grant #2047239.

## A. Appendix

### A.1 Table of parameters for simulation, and conditions for population trajectory and community dynamics

#### A.1.1 Simulation of two-species communities

We simulate 1500 two-species communities for each functional form by varying the parameters embedded in each function. We randomly drew 1500 sets of parameter values. We used each set of parameters to simulate the abundance of a two-species community for 500-time steps, starting at equal abundance. These parameters were randomly drawn from a distribution along their potential range.

- *N*_*o,ii*_,*N*_*o,ij*_,*N*_*o*, *jj*_, and *N*_*o,ii*_: the optimal density can theoretically have any positive value, yet we set the range between 0 and 10 and the initial density of 5.
- α_0,*ii*_ and α_0, *jj*_: the intraspecific density-independent effects were always negative as individuals of the same species are assumed to always compete at high densities. Yet, they can still facilitate themselves at low density depending on the density-dependent effect and the stretching parameter in the alternative functional forms.
- α_0,*ij*_ and α_0, *ji*_: the interspecific density-independent effects were drawn from a normal distribution centred around zero and with deviation of 0.2 as most interaction effects previously estimated for a Ricker model are usually below 0.1 (see Empirical example below). Interspecific density-independent effects were either exclusively positive (i.e., main interactive effects are facilitation), negative (i.e., main interactive effects are competition) or both (i.e., main interactive effects are competition and facilitation). We draw 250 sets of parameters specific to each of the three different scenarios of the main interactive effects
- α^_*ii*_, α^_*ij*_, α^_*jj*_ and α^_*ji*_: the inter- and intraspecific density-dependent effects defining the shape of the interaction were sampled from a uniform distribution from −1 to 0.
- *c*_*ii*_, *c*_*ij*_, *c* _*jj*_ and *c* _*ji*_: the stretching parameters were sampled from a uniform distribution from −1 to 0. The stretching parameters are negatives as α^ are negatives.

For the density-independent effect, we simulated 250 communities for each of the three different scenarios of main interactive effects: (i) interspecific effects are mainly positive over the density-gradient, both density-independent effects are positive (α_0,*ij*_ > 0 and α_0, *ji*_ > 0), (ii) interspecific effects are mainly competitive over the density-gradient, both density-independent effects are negative (α_0,*ij*_ < 0 and α_0, *ji*_ < 0) and, (iii) interspecific effects are in opposite direction, one mainly positive and one mainly negative, over the density-gradient, that is the density-independent effects have an opposite direction (α_0,*ij*_ > 0 and α_0, *ji*_ < 0, or inversely).

**Table S1:** Parameters used for simulating two-species community. Values of varying parameters were sampled within their range according to the specific distribution for each simulation.

| Parameters | Name | Range | Distribution for simulation |
| --- | --- | --- | --- |
| $\lambda_i$ and $\lambda_j$ | Intrinsic growth rate | constant | equal to 1 |
| $N_{o,ii}$, $N_{o,ij}$, $N_{o,jj}$, and $N_{o,ji}$ | Optimal density | [0,10] | $U(0, 10)$ |
| $\alpha_{0,ij}$ and $\alpha_{0,ji}$ | Interspecific density-independent effect | [-1,1] | $N(0, -0.2)$ |
| $\alpha_{0,ii}$ and $\alpha_{0,jj}$ | Intraspecific density-independent effect | [-1,0] | $N(0, -0.2)$ |
| $\hat{\alpha}_{ii}$, $\hat{\alpha}_{ij}$, $\hat{\alpha}_{jj}$ and $\hat{\alpha}_{ji}$ | Density-dependent effect | [-1,0] | $U(0, -1)$ |
| $c_{ii}$, $c_{ij}$, $c_{jj}$ and $c_{ji}$ | Stretching parameter | [-1,0] | $U(-1, 0)$ |

#### A.1.2 Population trajectory conditions

In one community, we characterised the populations’ abundance over time as either (i) both species persist, (ii) only one species persists, (iii) none persist or (iv) at least one species shows a run-away trajectory. In addition to evaluating the growth rate when low, we also categorised the population’s persistence by examining the median and maximum/minimum abundances reached.

Result in visualisation by tables.

**Table S2:** Categorisation of community trajectory and their respective conditions. GRWL= growth rate when low. N represent the abundance time series. The time series was considered in its entirety for detecting runaway dynamic (i.e., *N*_0,500_), but the 100 first-time steps were not considered for any other trajectories (i.e., *N*_100,500_).

| Community trajectory | Condition respected for both species | Condition |
| --- | --- | --- |
| Two species | Both | $\min(GRWL_{100,200}) > 1$ & $500 > \text{median}(N_{100,200}) > 0.05$ |
| One species | At least one | $\min(GRWL_{100,200}) > 1$ & $500 > \text{median}(N_{100,200}) > 0.05$ |
| No species | Both | $\text{median}(N_{100,200}) < 0.05$ or $GRWL < 0$ |
| Run-away | At least one | $\max(GRWL_{100,200}) > 100$ or $\max(N_{100,200}) > 500$ or $\text{median}(N_{0,500}) > 500$ |

#### A.1.3 Community trajectory conditions

Community synchrony describes the correlation of the populations’ abundance over time (Shoemaker et al., 2022). We can test for short-term and long-term synchrony by fitting the function *tsvreq*_c_ lassic from the package timescale-specific variance ratio version 1.0.2 (Zhao et al., 2024). The function *tsvreq*_c_ evaluates the presence of correlation between two timeseries (two series of abundances over time) that translate the presence of synchrony at long and/or short time scales (Zhao et al., 2020). Synchrony can be coupled with the oscillation of the populations over time. The oscillation was assessed by fitting an Autoregressive integrated moving average with the R function auto.arima() from the package *forecast version 8.21* (Hyndman et al., 2023). The function fits the best ARIMA model to a univariate time series and gives estimations of q (i.e., dimension of the moving average component) and p (i.e., dimension of the autoregressive component of the model) when needed (Ives et al., 2010). The evaluation of p and q tests the existence of oscillation dynamics, such as delayed density dependence.

**Table S3:** Summary of the community composition in the simulated experiment in percentage. In the first three columns (“Both compete”;“One competing,one facilitating”,“Both facilitate”), the percentage is the number of communities found to have that population outcome (e.g., runaway) with the predominant interactions specified by the columns, out of the 500 simulated. The “Total” is the number of communities found to have that population outcome across scenarios of predominant interactions, out of the 1500 simulated.

| functional form | population outcome | Both compete | One competing, one facilitating | Both facilitate | Total |
| --- | --- | --- | --- | --- | --- |
| Traditional constant | run away population | 0.2 | 1.6 | 76.4 | 26.2 |
| Traditional constant | no species | 0 | 0.4 | 0 | 0.13 |
| Traditional constant | one-species community | 84.8 | 59.4 | 1.6 | 48.85 |
| Traditional constant | two-species community | 15 | 38.6 | 22 | 25.33 |
| Linear | run away population | 1.4 | 0.8 | 0.6 | 0.94 |
| Linear | no species | 64.6 | 76.4 | 79.2 | 73.76 |
| Linear | one-species community | 15.6 | 9.4 | 7.6 | 10.93 |
| Linear | two-species community | 18.4 | 13.4 | 12.6 | 14.87 |
| Exponential | run away population | 2 | 10.4 | 29.8 | 14.13 |
| Exponential | no species | 54.8 | 51.6 | 42 | 49.72 |
| Exponential | one-species community | 23.6 | 15 | 4.6 | 14.48 |
| Exponential | two-species community | 19.6 | 23 | 23.6 | 22.18 |
| Sigmoid | run away population | 0.2 | 1.8 | 15 | 5.7 |
| Sigmoid | no species | 0.4 | 6.6 | 13.2 | 6.76 |
| Sigmoid | one-species community | 50.6 | 33.8 | 7 | 30.61 |
| Sigmoid | two-species community | 48.8 | 57.8 | 64.8 | 57.42 |

**Table S4:**
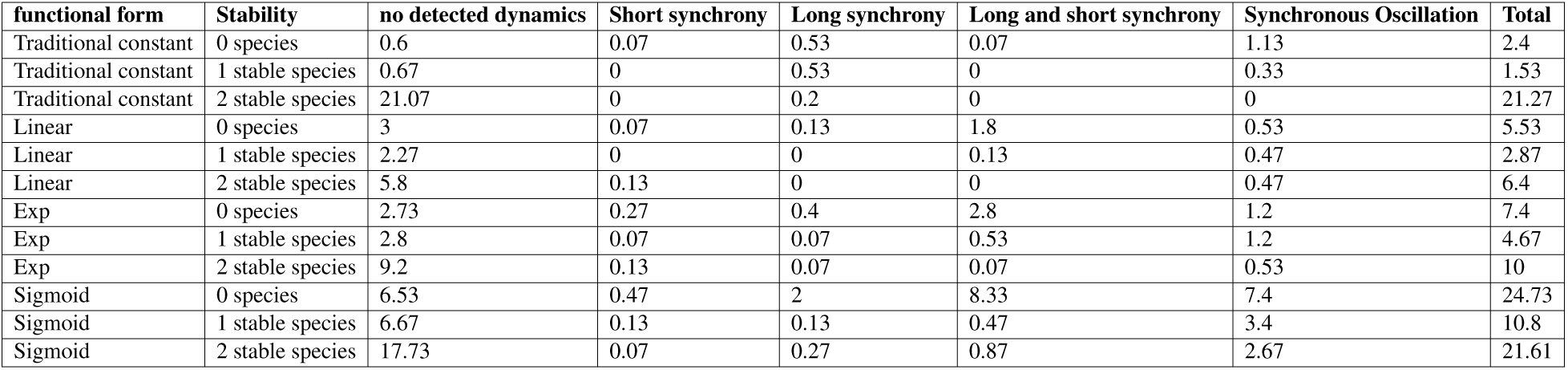
Summary of the community’s internal dynamics in the simulated experiment. Stability column describes the number of species detected as temporally stable.

| functional form | Stability | no detected dynamics | Short synchrony | Long synchrony | Long and short synchrony | Synchronous Oscillation | Total |
| --- | --- | --- | --- | --- | --- | --- | --- |
| Traditional constant | 0 species | 0.6 | 0.07 | 0.53 | 0.07 | 1.13 | 2.4 |
| Traditional constant | 1 stable species | 0.67 | 0 | 0.53 | 0 | 0.33 | 1.53 |
| Traditional constant | 2 stable species | 21.07 | 0 | 0.2 | 0 | 0 | 21.27 |
| Linear | 0 species | 3 | 0.07 | 0.13 | 1.8 | 0.53 | 5.53 |
| Linear | 1 stable species | 2.27 | 0 | 0 | 0.13 | 0.47 | 2.87 |
| Linear | 2 stable species | 5.8 | 0.13 | 0 | 0 | 0.47 | 6.4 |
| Exp | 0 species | 2.73 | 0.27 | 0.4 | 2.8 | 1.2 | 7.4 |
| Exp | 1 stable species | 2.8 | 0.07 | 0.07 | 0.53 | 1.2 | 4.67 |
| Exp | 2 stable species | 9.2 | 0.13 | 0.07 | 0.07 | 0.53 | 10 |
| Sigmoid | 0 species | 6.53 | 0.47 | 2 | 8.33 | 7.4 | 24.73 |
| Sigmoid | 1 stable species | 6.67 | 0.13 | 0.13 | 0.47 | 3.4 | 10.8 |
| Sigmoid | 2 stable species | 17.73 | 0.07 | 0.27 | 0.87 | 2.67 | 21.61 |

**Table S5:**
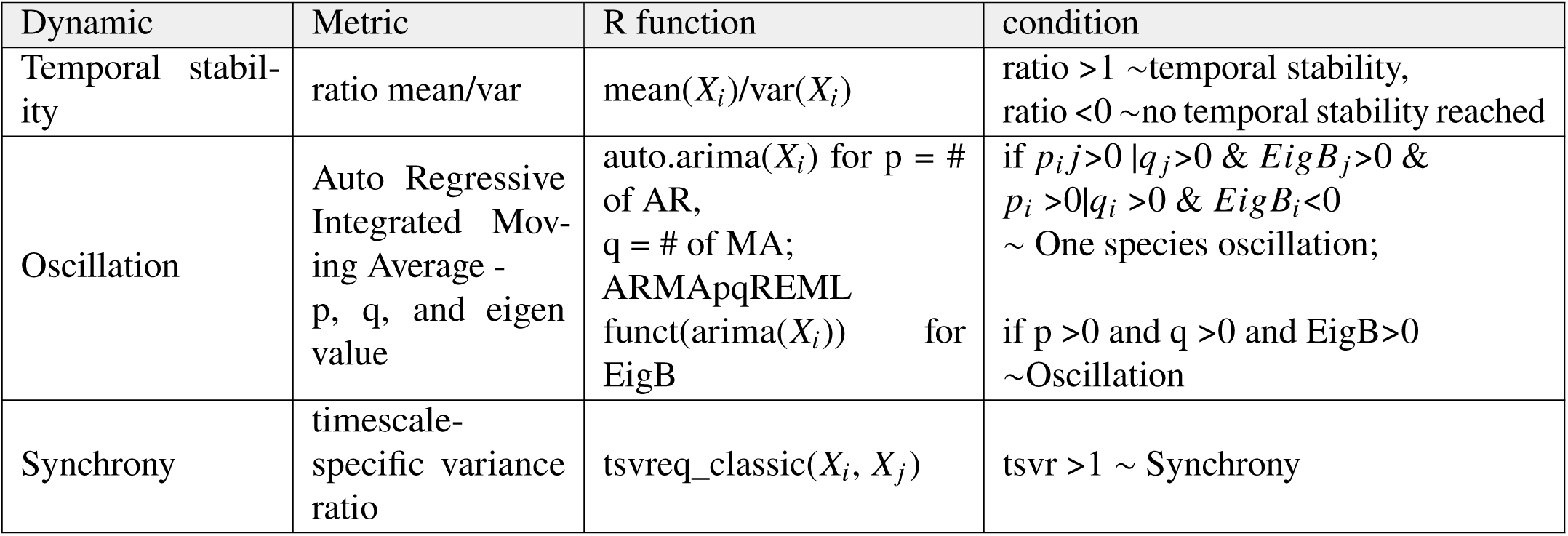
Table of metrics/criteria used to describe the dynamics and stability in each simulated community for each functional form. Temporal stability was assessed for each population in the community by looking at the ratio of the mean over the variance of the population’s density after 100 time-steps until the 200th. The Oscillation was assessed by fitting an Auto regressive integrated moving average with the R function auto.arima() from the package *forecast version 8.21* (Hyndman et al., 2023). The function fits the best ARIMA model to a univariate time series and gives estimations of q (i.e., dimension of the moving average component) and p (i.e., dimension of the autoregressive component of the model) when needed (Ives et al., 2010). The evaluation of p and q tests the existence of oscillation dynamics, such as delayed density dependence. Synchrony was assessed by fitting the function *tsvreq*_c_ from the package timescale-specific variance ratio version 1.0.2 (Zhao et al., 2024). The function evaluates the presence of correlation between two timeseries (two series of abundances over time) that translate the presence of synchrony at long and short time scales (Zhao et al., 2020).

### A.2 External conditions

To highlight the importance of facilitation coupled with external factors, we run all analyses under three different scenarios of external factors. Indeed, a change in the abiotic environment can drive species to low abundance and ultimately push them to extinction. Facilitation can rescue species and change our prediction of which species can persist in parallel with environmental variation.

We added an extra parameter to the descriptions of *N*_*i*,t_, the density of species *i* at time t. We used the exponential discrete Lotka-Volterra, also called a Ricker model (Ricker, 1954) with the addition of an eternal pressure *E*:

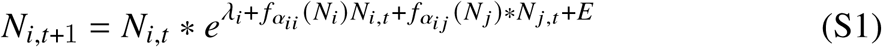

with γ_*i*_ the intrinsic growth rate, *f*_α*ij*_ the functional form of the effect if *j* on *i, E* is an external pressure described in three ways (i) non-existent *E* 0;(ii) a periodic perturbation defined by sine function *E sin*(2π/*b*) or (iii) a pulse perturbation defined by a random sample function *E N* (0, *b*). The parameter *b* is specific for each community and equal for both species, sampled from a uniform distribution *b U*(0, 2). We visualise the three scenarios for one set of parameters in Fig.**??**.

The positive correlations emerging from facilitation are expected to diminish community robustness against stochastic events (). To test how the inclusion of varying species interactions impacts that hypothesis, we run all analyses under three different scenarios of external factor: (i) non-present (“No external factor”, presented in the main text), (ii) stochastic (“Noisy change”, i.e. *E N* (0, *b*)) and, (iii) periodic (“Periodic change”, i.e., *E sin*(2π/*b*)) with *b* specific for a community and equal for both species.

**Figure S1:**
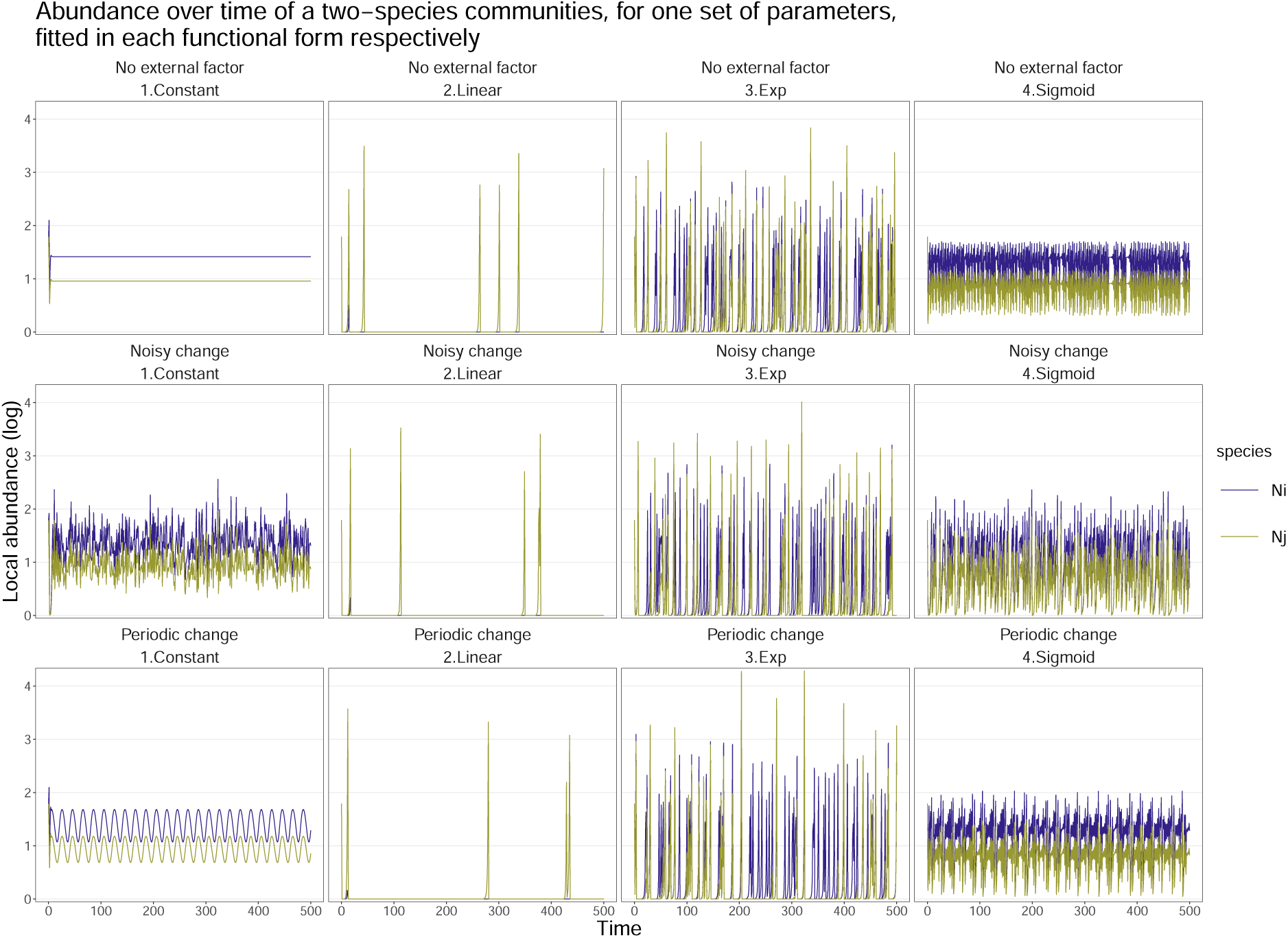
The four density-dependent functional forms present different local community dynamics. The community is defined by the same set of parameters (i.e., for interaction *i j*: α_0,*ij*_,α_*ij*_,*N*_0_,α^^^_*ij*_). The community dynamics is described by a Ricker model, eq.7, with fecundity defined by eq.6. An external factor acts on species’s fecundity. The external factor (represented by the black line) is either non-present (“No external factor”), stochastic (“Noisy change”, i.e. *E N* (0, *b*)) or periodic (“Peridoc change”, i.e., *E sin*(2π/*b*)) with *b* specific for this community and equal for both species. We represent 50 years out of the 100 simulated, for simulation number 100, out of the 500 simulated for each scenario.

**Figure S2:**
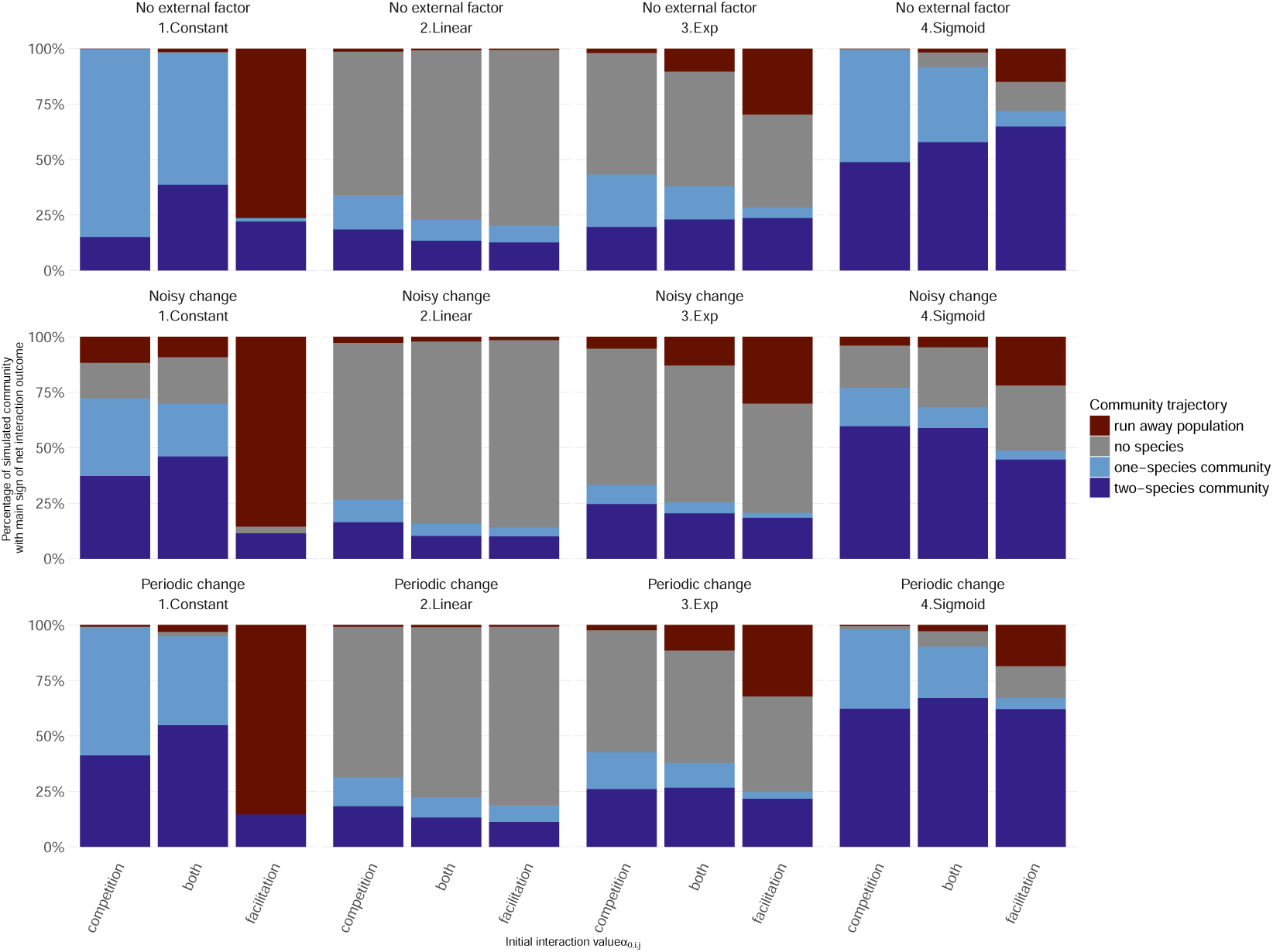
Community trajectory differs based on the functional form used and the governing interactions. Under main facilitation, the traditional constant form shows mainly runaway dynamics, contrary to all other functional forms. The linear and exponential linear forms include an impressively high number of “no species” communities. The sigmoid form shows a majority of high diversity communities, regardless of the governing interactions. The community trajectory is based on the minimum growth rate at low abundance (>0). Runaway dynamics is reached when density exceeds 1000 individuals, corresponding to the initial density times 1000. The community trajectory is evaluated based on three initial conditions/abundance of the two species: (ii) equal invasion (*N*_*i*_=*N* _*j*_ =5), (i) Species *i* invades species *j* at its equilibrium (*N*_*i*_=1, *N* _*j*_ =*N*_*j*_^∗^),(i) inversely (*N*_*i*_=*N*_*j*_^∗^, *N*_*j*_ =1). The minimum growth rate is determined based on the three communities.

**Figure S3:**
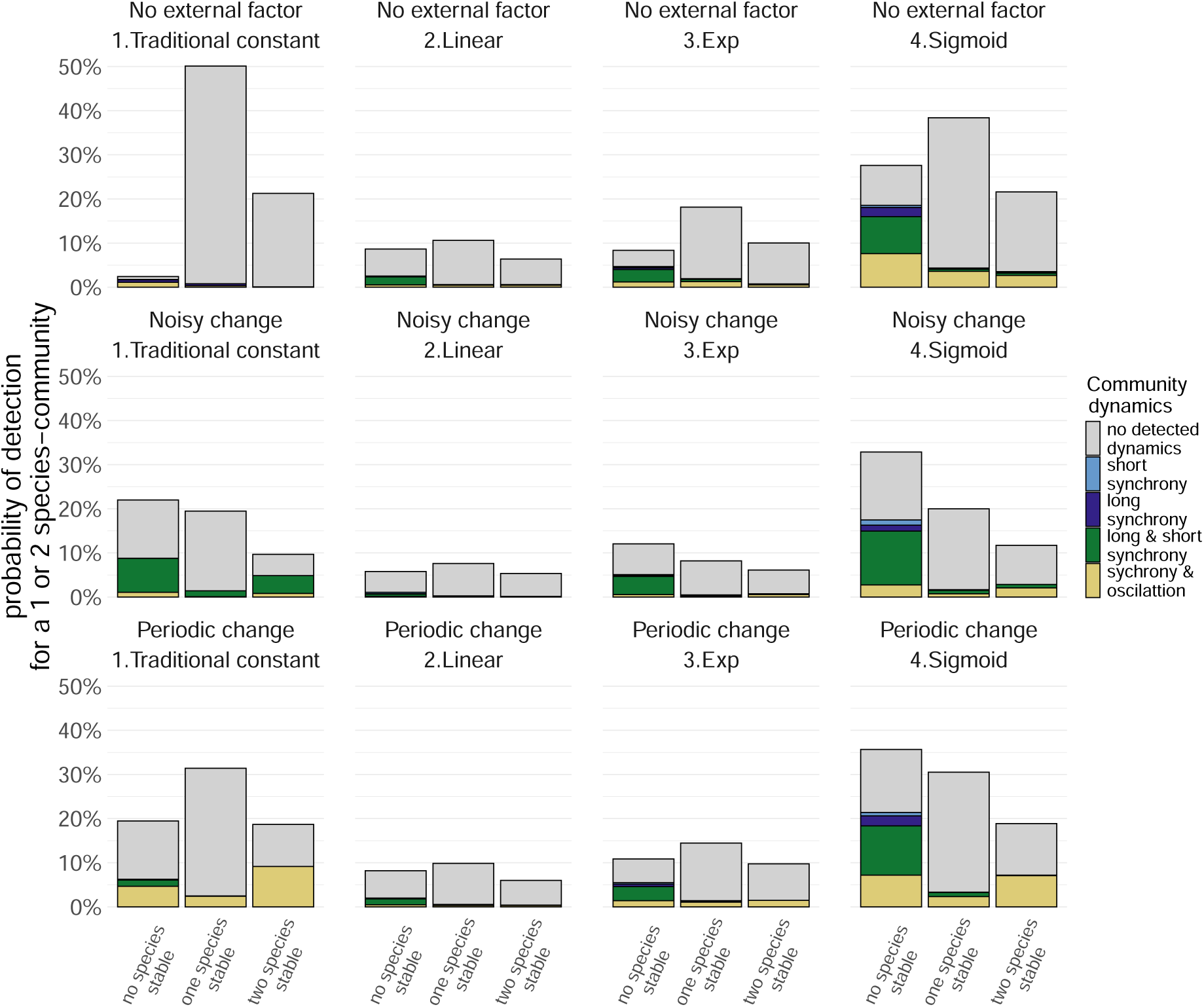
The different functional forms can lead to diverse community dynamics, yet the sigmoid form is the most versatile. The number of communities predicted to have at least one species persisting is also the highest in the sigmoid form, followed by the constant form. We classified each community according to its dynamics, respectively, for each functional form, following Tab.S5. The bar plot level shows the total number of communities predicted to have at least one persisting species out of the 1500 simulated for each functional form.

The inclusion of external factors, in parallel to varying functional forms of species interactions, shows how facilitation can rescue species under external factors (here noisy/stochastic, periodic such as seasonal change, or punctual) and change our predictions as to which species will persist in parallel with environmental variations. Indeed, two non-coexisting species do not instantly exclude one another. For instance, an annual plant can take 15/20 years to be excluded or more than hundreds of years for perennial plants (Chu and Adler, 2015). The exclusion of a species from a community is a long pending process during which complex dynamics (switch of species interactions, introduction of a new species, environmental stochasticity, anthropogenic perturbations, nutrient level) can modify the outcome of species interactions and population trajectory (e.g., DeSiervo et al., 2023).

Concerning the population trajectory, we see more species predicted to not persist when introducing noisy change in per capita growth rate, but similar result between the absence of external factors and periodic change (Fig.S2). For the community dynamics, we see a reduction in stable population across all functional forms (Fig.S3). Yet, this reduction is more drastic for the constant function than its “no external factor scenario”. Overall, the conclusion of the importance of the inclusion of a varying bounded functional form is maintained under all scenarios Fig.S2 and Fig.S3.

### A.3 IGR with varying species interactions

Despite the traditional use of the invasion growth rate (IGR) 1, it is impossible to use this standard metric due to its assumptions of a stationary environment, where the invader growth rate follows a constant value averaged over a long-time period (Barabás et al., 2018). Indeed, despite their ability to invade, species can be driven to extinction, and inversely, due to their lack of invasion probability shadows their ability to coexist in other conditions (Barabás et al., 2018). This mismatch of prediction and trajectory is due to the presence of more complex population dynamics. Moreover, the IGR assumes the invasion of one species within the stable community (Johnson and Hastings, 2022). That is, the community has to reach a stationary state before introducing the invader species. Yet, if the community relies on the invaders to reach a stable state (e.g., rock-paper-scissor, no two-species communities can stably coexist without the third species), such assumptions can not be made. These assumptions hinder the application of IGR with non-traditional functional forms (linear, exp, and sigmoid) and in natural scenarios of joint invasion.

### A.4 Exploration of parameters

Exploration of prediction of fig.8. The alternative functional forms of species interaction introduce new parameters which allow to capture more natural complexity. We evaluate the impact of these parameters on the community’s temporal stability. That is, we draw the linear relationship (glm, package *stats*) between the parameters within each functional form and the probability for temporal stability (ratio of mean over variance over time). For instance, for the linear functional form, we tested the following model:

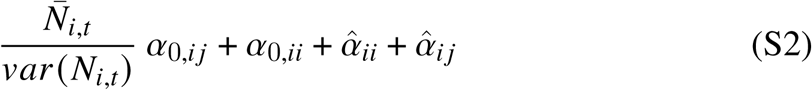

For each model, we can quantify the effect of each parameter on temporal stability (Tab.S6). We can then predict the relationship between each parameter and temporal stability (Fig. S3). The results extend the previous conclusion made in box.5.

**Table S6:** Effect of each parameter on the probability of the community to be stable (1= stable community, 0 = non-stable community) within a community predicted to coexist. Each functional form was fitted separately into a (glm, package *stats*), and the coefficients of each parameter and their interactions were estimated.

| funct.form | parameters | Estimate | Std. Error | t value | Pr(>\|t\|) |
| --- | --- | --- | --- | --- | --- |
| 1.constant | (Intercept) | -0.477 | 0.092 | -5.169 | 0.000 |
| 1.constant | alpha.0.intra | -0.677 | 0.090 | -7.558 | 0.000 |
| 1.constant | alpha.0.inter | 0.423 | 0.096 | 4.412 | 0.000 |
| 1.constant | alpha.0.intra:alpha.0.inter | 0.002 | 0.091 | 0.026 | 0.980 |
| 2.linear | (Intercept) | -1.380 | 0.151 | -9.162 | 0.000 |
| 2.linear | alpha.0.intra | -0.025 | 0.126 | -0.197 | 0.844 |
| 2.linear | alpha.0.inter | 0.212 | 0.150 | 1.416 | 0.158 |
| 2.linear | alpha.hat.intra | 0.162 | 0.140 | 1.151 | 0.251 |
| 2.linear | alpha.hat.inter | -0.117 | 0.141 | -0.828 | 0.409 |
| 2.linear | alpha.0.intra:alpha.0.inter | 0.114 | 0.128 | 0.894 | 0.373 |
| 2.linear | alpha.hat.intra:alpha.hat.inter | -0.066 | 0.132 | -0.498 | 0.619 |
| 3.exp | (Intercept) | -1.737 | 0.128 | -13.545 | 0.000 |
| 3.exp | alpha.0.intra | -0.147 | 0.115 | -1.273 | 0.204 |
| 3.exp | alpha.0.inter | 0.125 | 0.112 | 1.124 | 0.262 |
| 3.exp | alpha.hat.intra | 0.025 | 0.119 | 0.206 | 0.837 |
| 3.exp | alpha.hat.inter | -0.327 | 0.122 | -2.687 | 0.008 |
| 3.exp | C.intra | 0.105 | 0.117 | 0.899 | 0.369 |
| 3.exp | C.inter | 0.102 | 0.115 | 0.890 | 0.374 |
| 3.exp | N.opt.intra | -0.164 | 0.125 | -1.310 | 0.191 |
| 3.exp | N.opt.inter | -0.018 | 0.127 | -0.140 | 0.888 |
| 3.exp | alpha.0.intra:alpha.0.inter | -0.086 | 0.111 | -0.778 | 0.437 |
| 3.exp | alpha.hat.intra:alpha.hat.inter | -0.195 | 0.119 | -1.638 | 0.102 |
| 3.exp | C.intra:C.inter | 0.012 | 0.118 | 0.105 | 0.917 |
| 3.exp | N.opt.intra:N.opt.inter | -0.012 | 0.124 | -0.097 | 0.923 |
| 4.sigmoid | (Intercept) | -1.490 | 0.065 | -23.007 | 0.000 |
| 4.sigmoid | alpha.0.intra | -0.544 | 0.065 | -8.333 | 0.000 |
| 4.sigmoid | alpha.0.inter | 0.117 | 0.064 | 1.819 | 0.069 |
| 4.sigmoid | alpha.hat.intra | 0.218 | 0.064 | 3.422 | 0.001 |
| 4.sigmoid | alpha.hat.inter | -0.061 | 0.066 | -0.930 | 0.353 |
| 4.sigmoid | C.intra | 0.284 | 0.065 | 4.380 | 0.000 |
| 4.sigmoid | C.inter | 0.029 | 0.064 | 0.458 | 0.647 |
| 4.sigmoid | N.opt.intra | -0.207 | 0.064 | -3.228 | 0.001 |
| 4.sigmoid | N.opt.inter | 0.031 | 0.065 | 0.487 | 0.626 |
| 4.sigmoid | alpha.0.intra:alpha.0.inter | 0.108 | 0.063 | 1.729 | 0.084 |
| 4.sigmoid | alpha.hat.intra:alpha.hat.inter | -0.091 | 0.064 | -1.428 | 0.154 |
| 4.sigmoid | C.intra:C.inter | 0.038 | 0.064 | 0.598 | 0.550 |
| 4.sigmoid | N.opt.intra:N.opt.inter | -0.035 | 0.064 | -0.544 | 0.586 |

**Figure S4:**
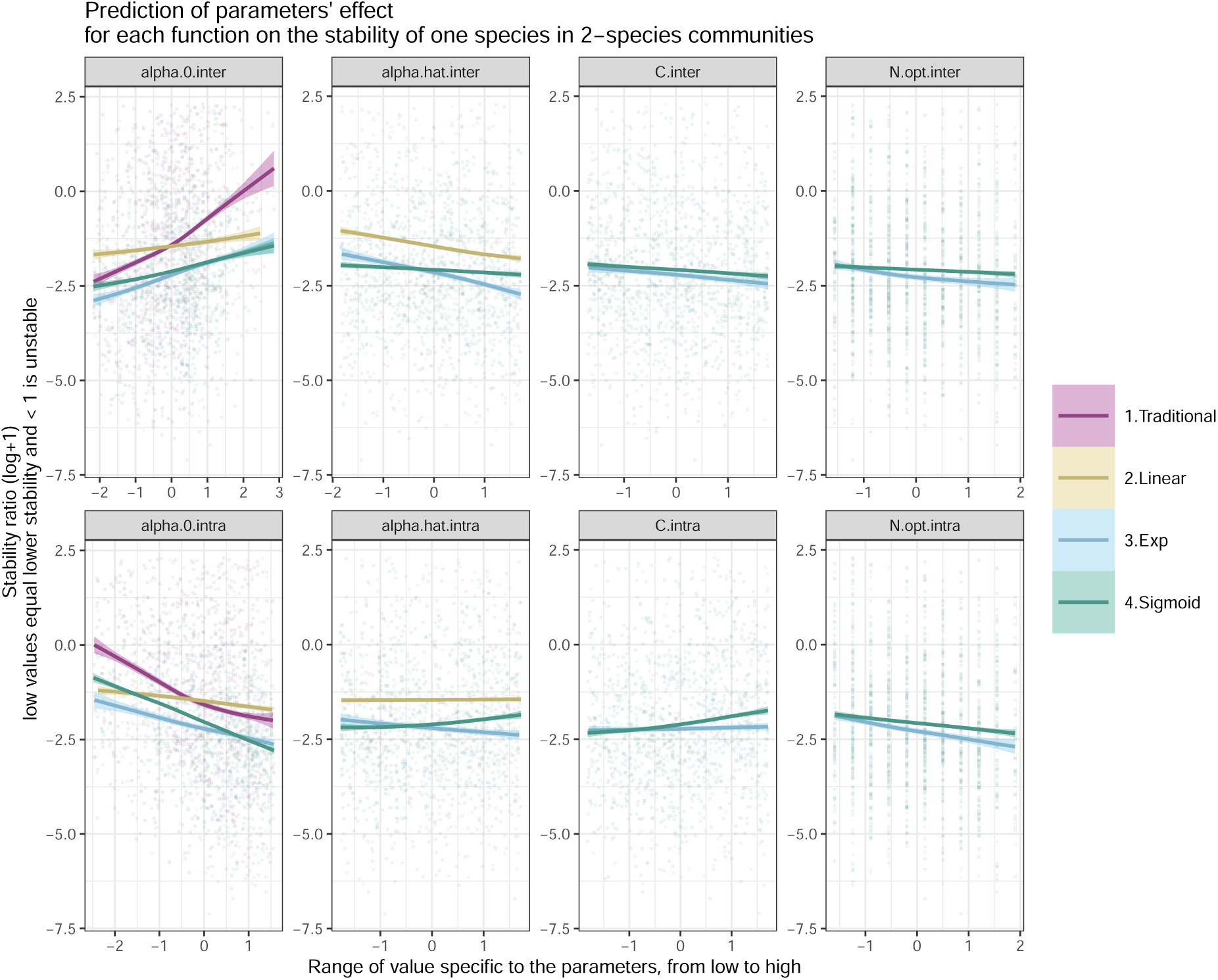
The prediction of reaching stability based on the function used. Only the linear function has, in majority, unstable communities.

**Figure S5:**
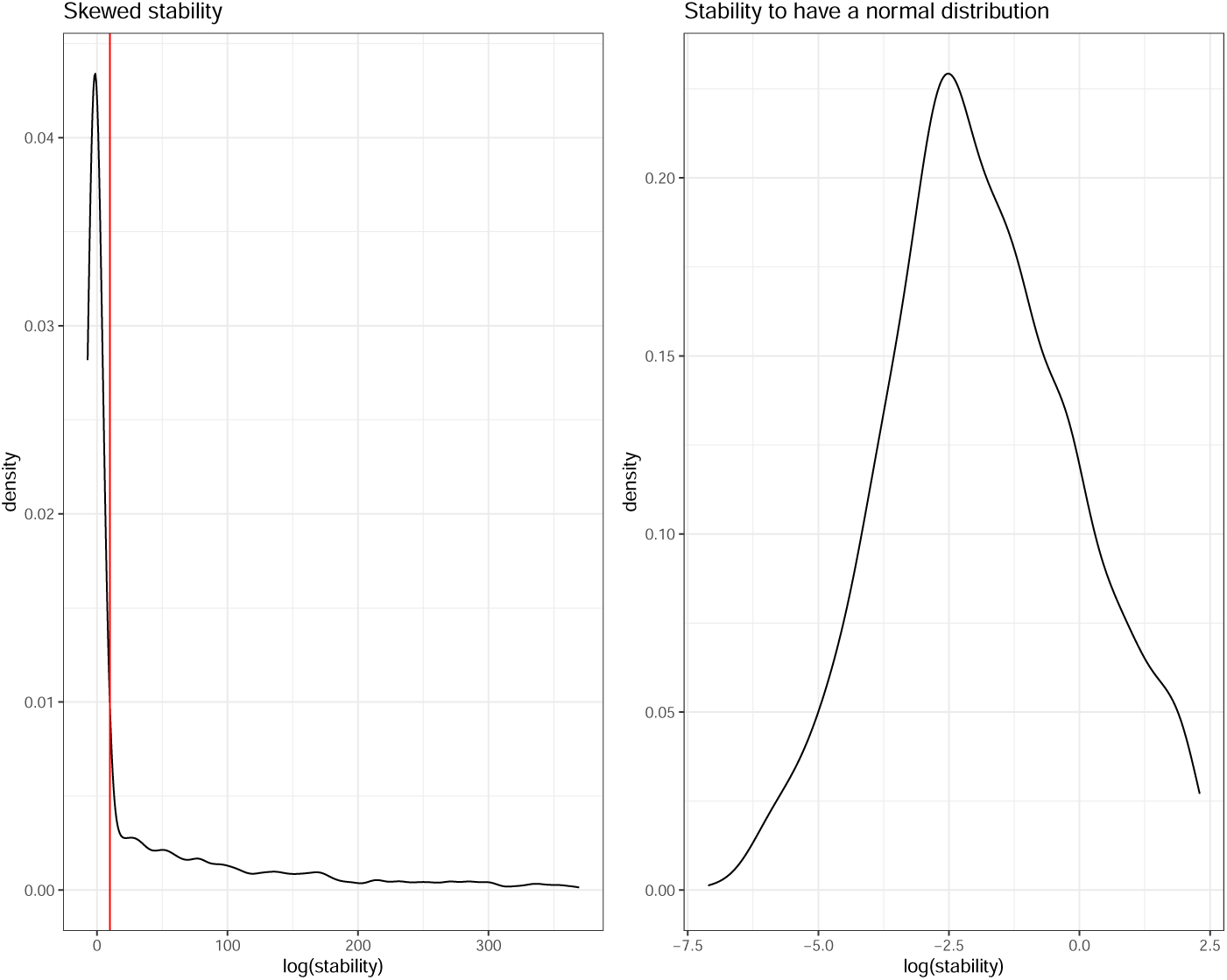
Stability ratio distribution when logged, with and without threshold. The values of the ratio between [-7,7] were kept to fit the glm. The thresholds consider realistic stability ratios and focus on how parameter values explain the switch from non-stable (ratio below zero) to stable (ratio above zero).

**Figure S6:**
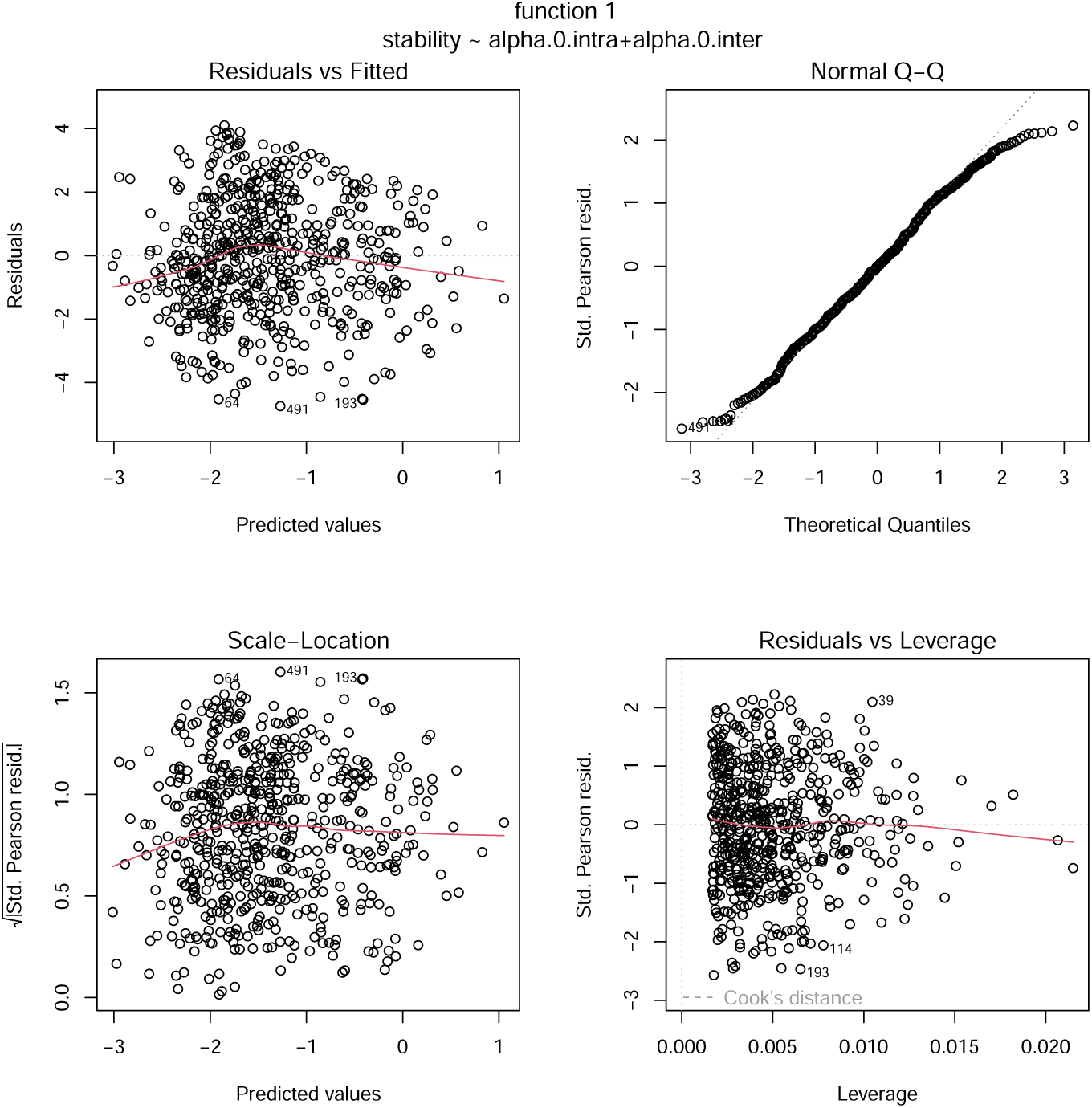
Diagnostic plots of the generalised linear model fitted to stability ratio as a response variable and independent effects as explanatory variables, specific to functional form 1.

**Figure S7:**
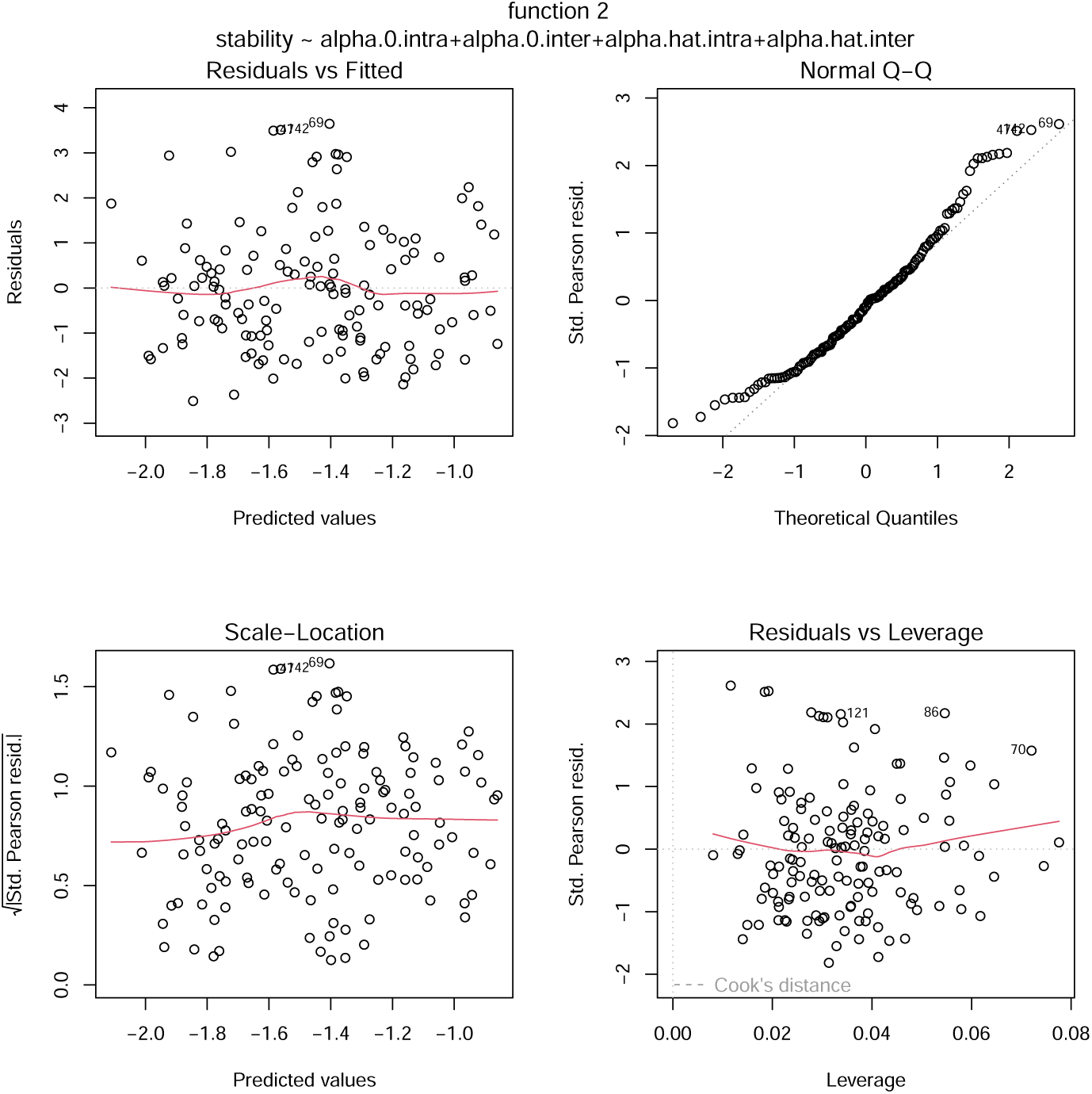
Diagnostic plots of the generalised linear model fitted to stability ratio as a response variable and parameter specific to functional form 2 as explanatory variables.

**Figure S8:**
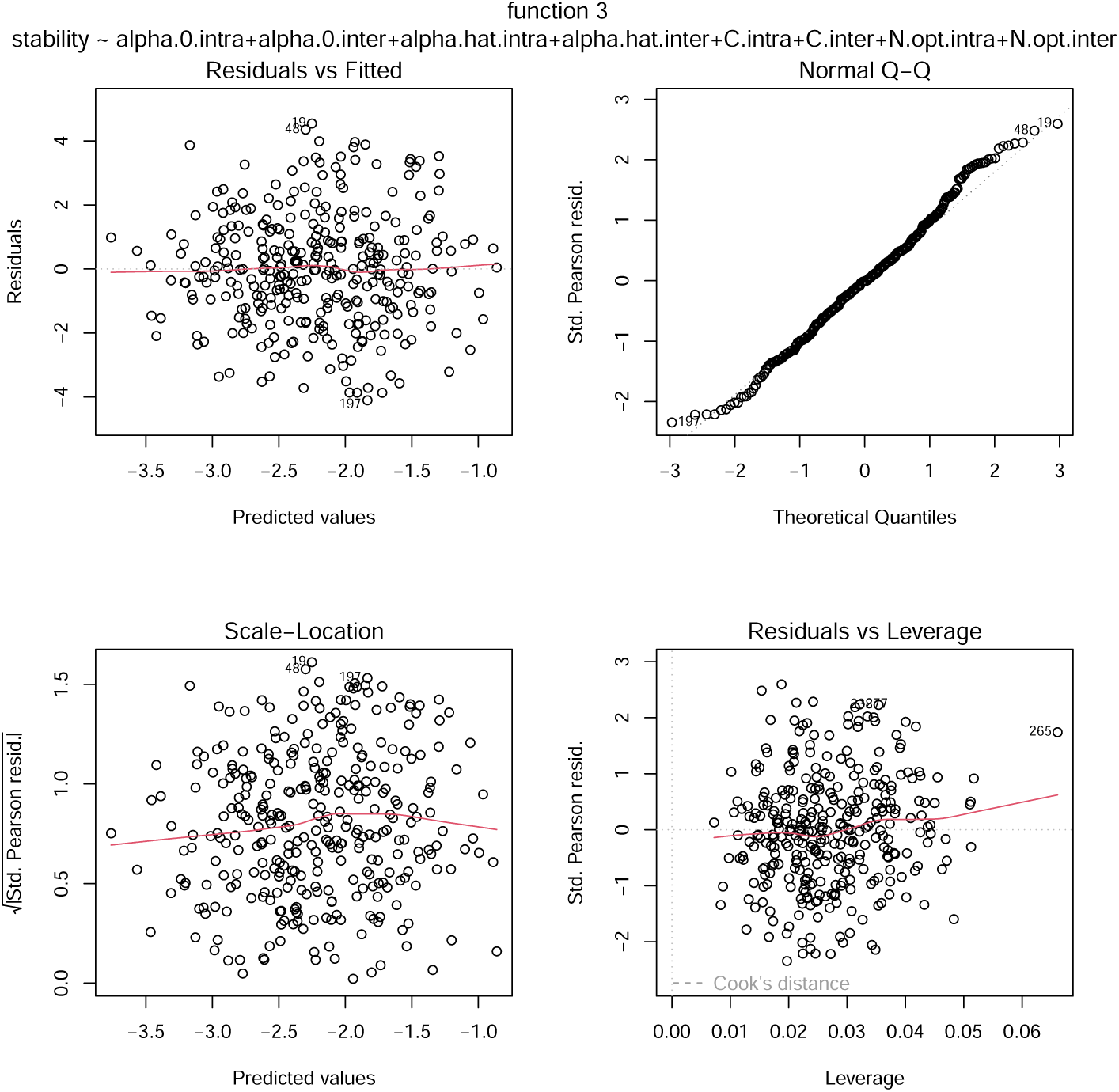
Diagnostic plots of the generalised linear model fitted to stability ratio as a response variable and parameter specific to functional form 3 as explanatory variables.

**Figure S9:**
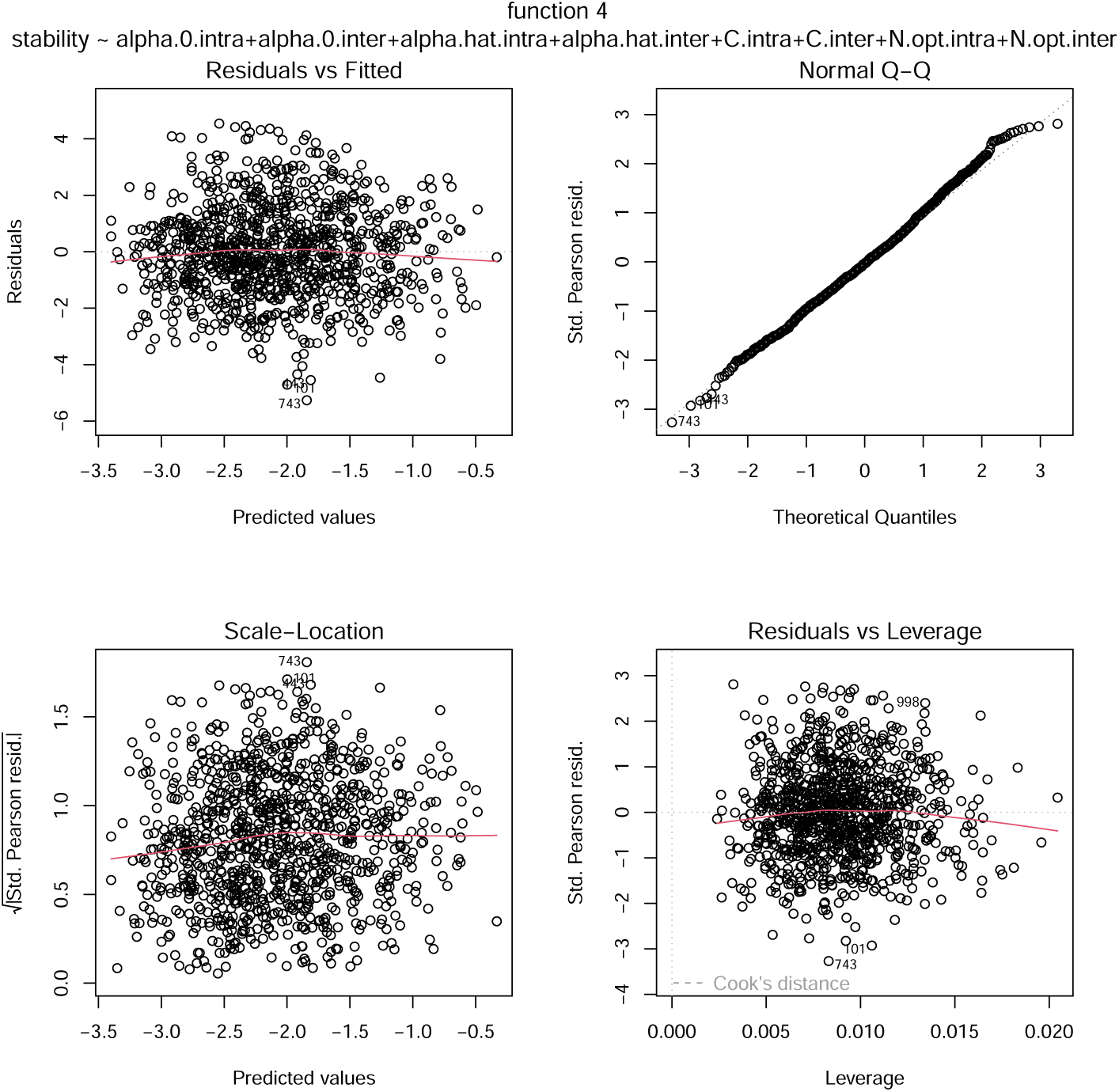
Diagnostic plots of the generalised linear model fitted to stability ratio as a response variable and parameter specific to functional form 4 as explanatory variables.

**Table S7:** Summary of the projection of the LARO population under each functional form.

| functional form | min | max | mean | standard deviation | median | Number of point |
| --- | --- | --- | --- | --- | --- | --- |
| Observed | 0 | 40 | 1.84 | 3.68 | 0.5 | 815 |
| 1.Traditional | 0 | 66.418 | 10.394 | 8.113 | 8.560 | 3000 |
| 2.Linear | 0 | 28.125 | 0.790 | 2.491 | 0.002 | 3000 |
| 3.Exp | 0 | 34.528 | 5.321 | 5.698 | 3.368 | 3000 |
| 4.Sigmoid | 0 | 41.476 | 6.669 | 6.480 | 4.640 | 3000 |

### A.5 Natural Data

#### A.5.1 Extended results

#### A.5.2 Why a Ricker model?

In the case of discrete counts of a performance metric, such as annual plant fecundity, community ecologists have frequently used the Beverton-Holt model (Watkinson, 1980) and the Ricker model (Ricker, 1954), which are both a discrete analogue of the Lotka-Voltera. While both models have been frequently used (e.g., (Godoy and Levine, 2014; Godoy et al., 2014; Bimler et al., 2018), the selection of an individual performance model form ultimately relies on the relevance for focal processes such as the performance’s metric distribution.

#### A.5.3 Bayesian model

Priors of parameters from eq.6 were set to follow ∼ *N* (0, 1)^−^. The density-dependent parameters (α^) and the stretching parameters (α^^^) had distribution restricted to [−0.5, 0.5] and later scaled by −0.5 to have value only between [−1, 0]. This operation led to higher values of both parameters and better convergence of their respective chains. The intrinsic growth rate (γ) had an upper limit defined by the highest observation of fecundity of LARO across all sampling. We limit the intrinsic growth rate to ensure it has realistic values rather than allowing it to adopt highly unrealistic values, which would lead to the overestimation of negative effects in species interactions. Similarly, the neutral neighbour densities were set here by the neighbour conditions concordant with the highest fecundity observed. For instance, we set the neutral neighbour density of LARO to 1 if we observed one individual of LARO in the neighbour of the focal individuals with the highest number of seeds produced.

#### A.5.4 Poisson multivariate regression

**Figure S10:**
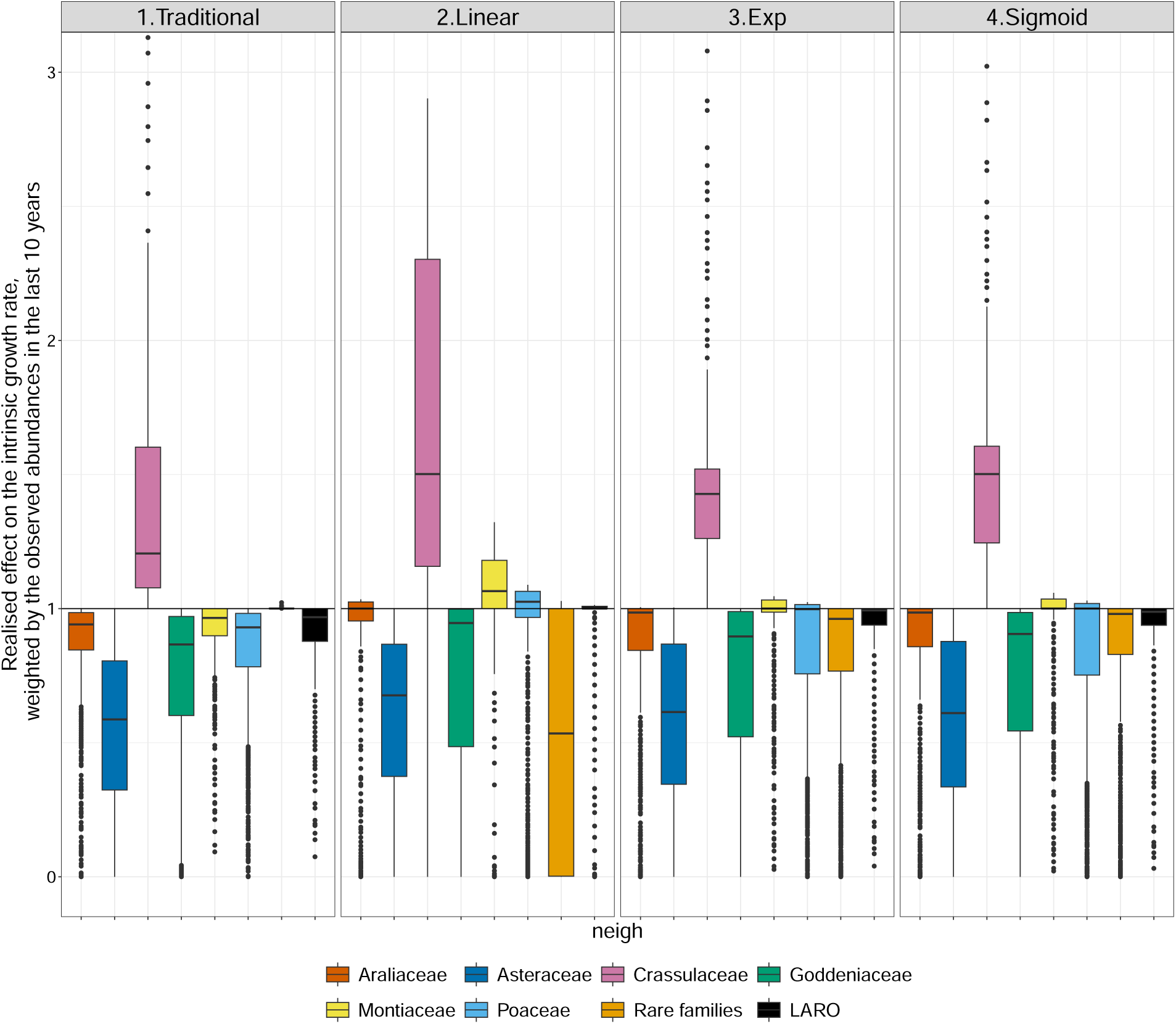
Realised effect of neighbours on LARO’s intrinsic fecundity, effect above one are increasing LARO’s fecundity (i.e., facilitation) and inversely for effect below one (i.e., competition). The effects are weighted by each family’s observed abundance in the last 10 years. For instance, the Araliaceae family, when present in LARO’s local neighbourhood, decreases its fecundity by 5% (average around 0.95) according to the traditionally constant function. LARO also experienced the most positive effect from the Crassulaceae and the most competitive effect from the Asteraceae family, which is the most abundant family of the system. Yet, we see differences between the functional forms, such as, when using one of the alternative functional forms (i.e., linear, exponential or sigmoid), individuals of the rare families negatively impact LARO, and the Montiaceae and Poaceae families positively impact LARO.

**Figure S11:**
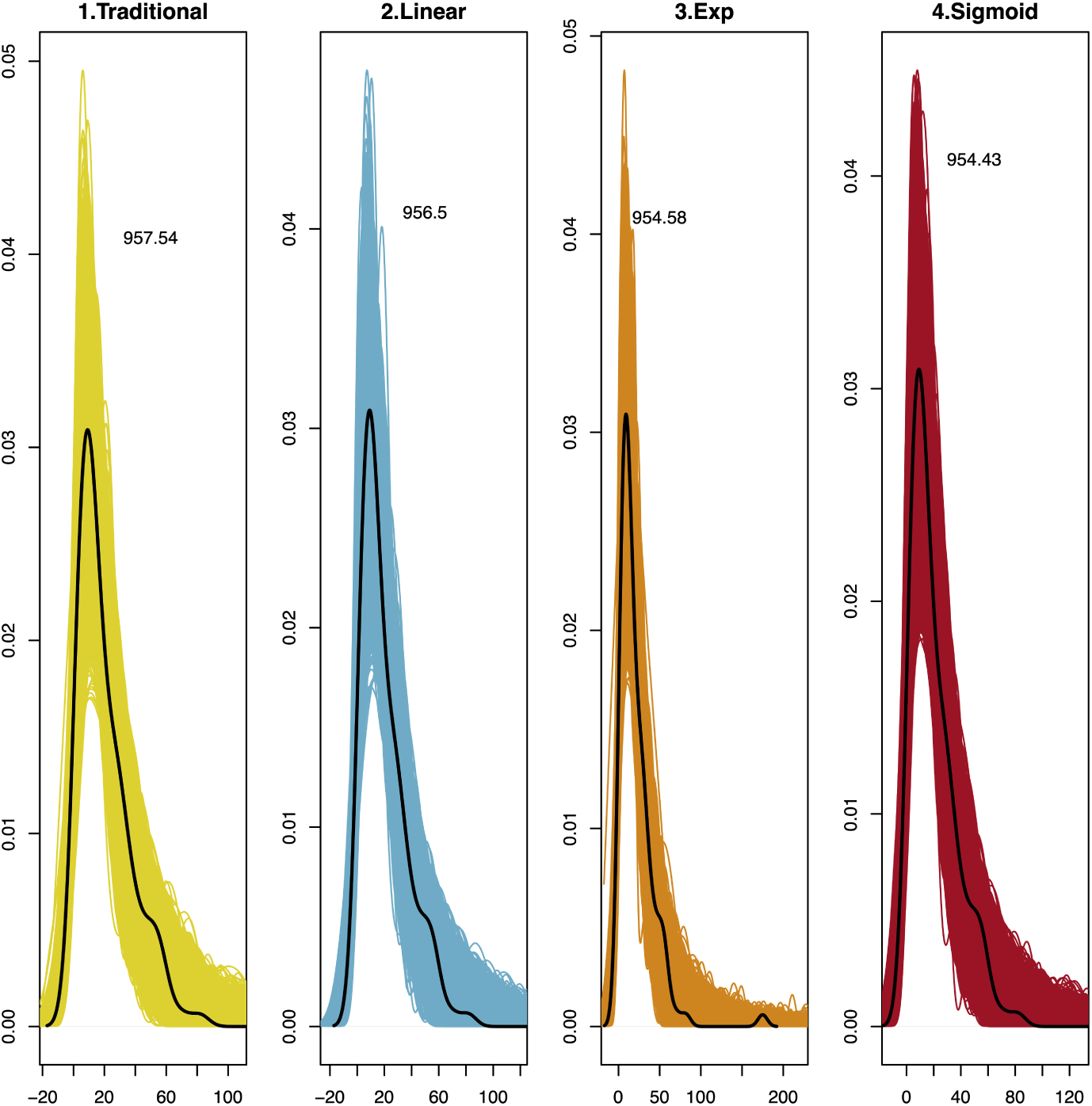
Posterior distribution of LARO’s fecundity. The black line shows the observed data. The colored lines show the posterior distribution for each iteration.

**Figure S12:**
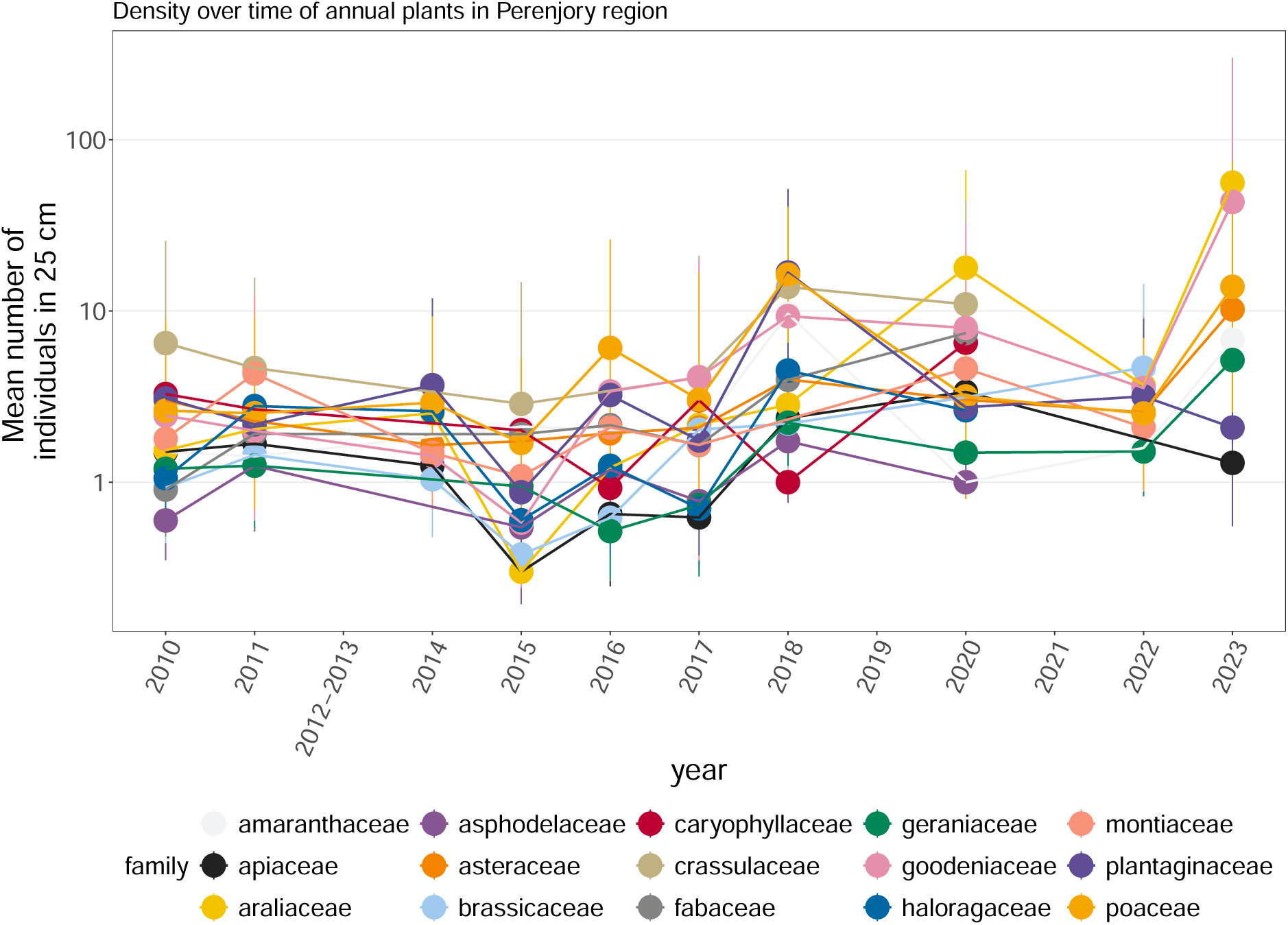
Time series of the density of the main families in Perenjori, WA, Australia, in 15 cm plot.

**Table S8:** Model behaviour when fitting the 4 different functional forms of species interactions for the natural community. Indicator: (1) Convergence using the Gelman–Rubin diagnostic (Rhat):*Rhat* < 1.1; Precision of parameter estimates using the effective sample size (Neff): *N Neff* > 100; (3) effective number of parameters (*P* − *loo*); (4) the theoretical expected log pointwise predictive density for a new dataset (*elpd*_*loo*_ in the “loo” R package version 2.7.0 Vehtari et al., 2024). We use the *loo*_*c*_*ompare* function (*loo*_*c*_*ompare* in the “loo” R package version 2.7.0 Vehtari et al., 2024), which compares the *elpd*_*loo*_ (i.e., log probability of the ability of the model to predict a new dataset). From *elpd*_*loo*_, we can compare multiple model fits of one dataset (here between different groups of the neighbourhood: class, family or species) by comparing the *elpd*_*diff*_. *SE*_*diff*_ is the standard error of component-wise differences of *elpd*_*loo*_ for two models. If *SE*_*diff*_ ∗ 1.96 < than *elpddiff*, then the models are potentially significantly different (Vehtari et al., 2017). Yet we need to also compare their LOOIC which can be used similarly to an AIC, where smaller values indicate more parsimonous models. We see here that while the sigmoid model has the lowest LOOIC, and the tradition model the highest, the values between models overlap greatly when considering the standard errors.

| function | N obs | Rhat | Neff | $elpd_{diff}$ | $se_{diff}$ | $elpd_{loo}$ | $se_{elpd_{loo}}$ | $p_{loo}$ | $se_{p_{loo}}$ | looic | $se_{looic}$ |
| --- | --- | --- | --- | --- | --- | --- | --- | --- | --- | --- | --- |
| 1.traditional | 118 | 1.002 | 2379.236 | -1.556 | 1.160 | -478.769 | 10.690 | 7.701 | 1.532 | 957.538 | 21.379 |
| 2.linear | 118 | 1.002 | 1987.188 | -1.039 | 1.430 | -478.252 | 9.887 | 9.208 | 1.652 | 956.504 | 19.775 |
| 3.exp | 118 | 1.002 | 1689.244 | -0.078 | 0.340 | -477.291 | 10.138 | 8.230 | 1.570 | 954.583 | 20.277 |
| 4.sigmoid | 118 | 1.003 | 1106.157 | 0.000 | 0.000 | -477.213 | 10.233 | 7.800 | 1.386 | 954.426 | 20.466 |

**Figure S13:**
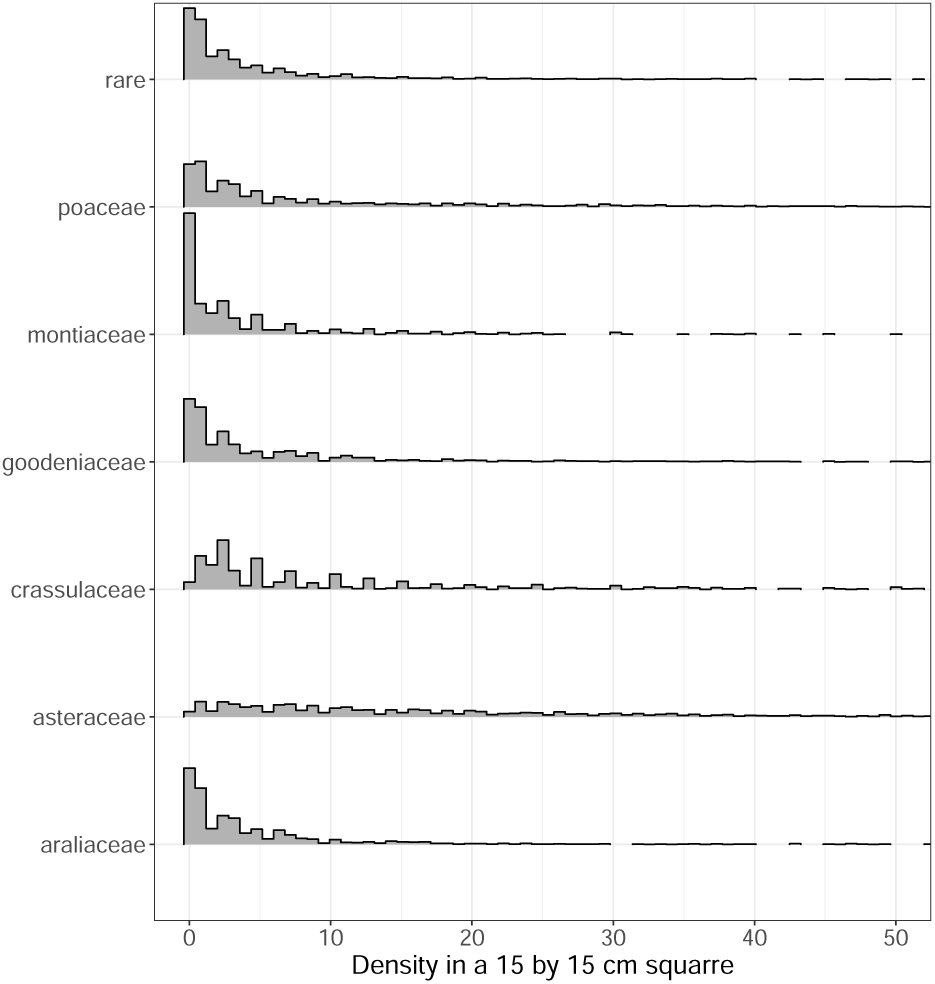
We can see the observed distribution of each family’s abundance used to fit the multivariate poisson.

**Figure S14:**
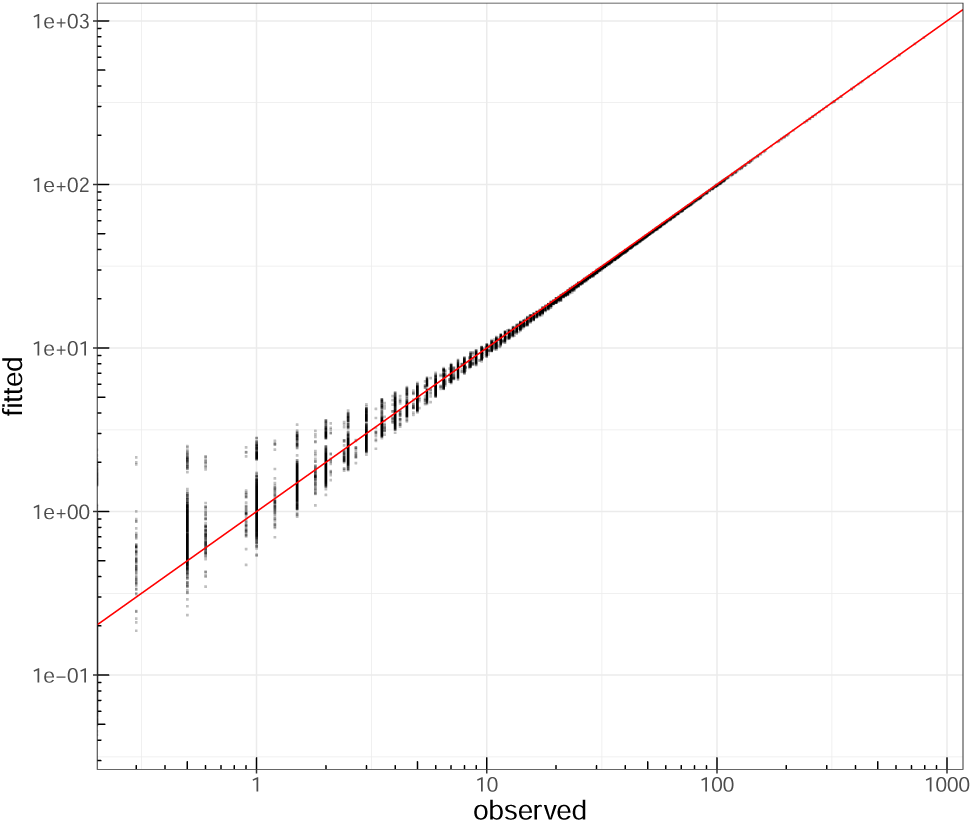
To check that the algorithm learned correctly from the data, we can look at the fitted value of the counts vs. observation. This indicates good performance overall, yet, at low abundances the model tends to overpredict.

**Figure S15:**
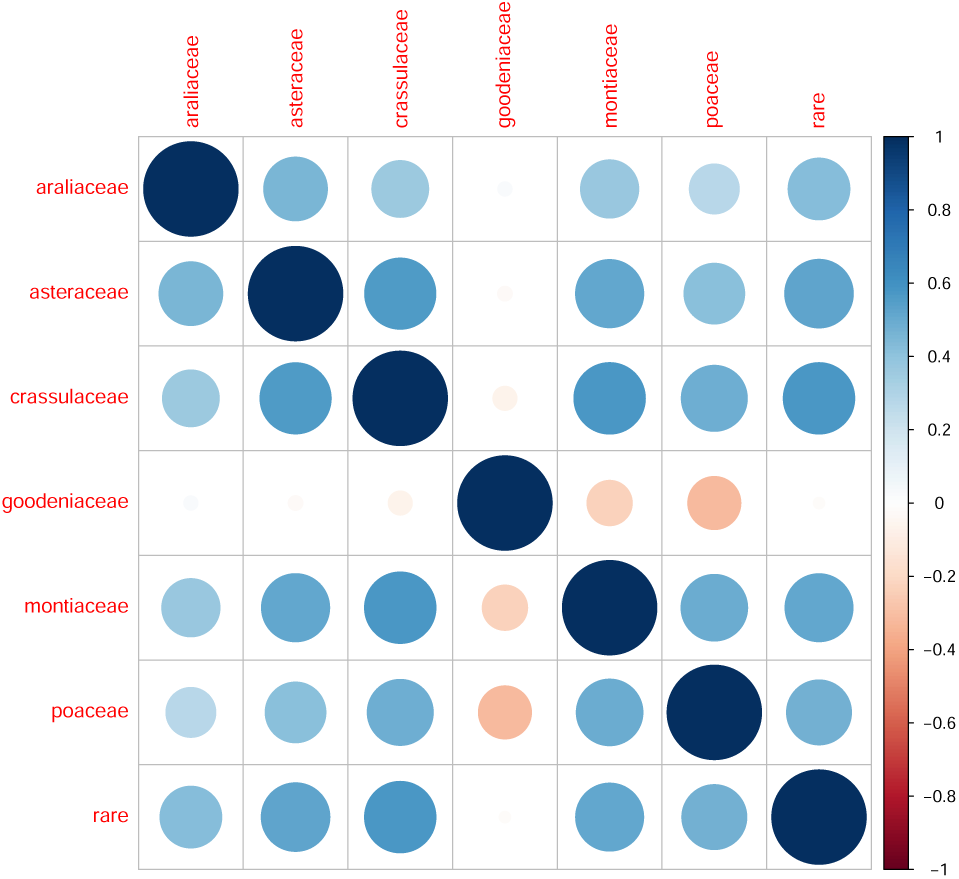
We computed the residuals correlation matrix between each family’s density across the 13 years by fitting a multivariate Poisson to our observed time-series (PNL, *PLNmodels* package Chiquet et al., 2021; appendix, Fig.S15). The multivariate Poisson accounts for correlation patterns in density and allows us to maintain these patterns in the simulated time series (appendix, Fig.S13,S14).

### A.6 Recovering parameters

We evaluated the capacity of the functional forms to recover predefined parameters. We sampled a list of 100 parameter sets and created for each set a dataset of a surface experiment with two competing species (abundance of 2 species and their performance). The parameters were sampled according to the following distribution:

- germination and seed survival were constant for both species and set to 0.9
- “lambda” is the intrinsic fecundity; for each dataset, two values (one per focal) were sampled as γ ∼ [20, 10] within the range of [10, 30].
- “alpha-init” is the constant density-independent effect; for each dataset, 4 values were sampled as α_0_ ∼ [−0.05, 0.1] within the range of [−0.5, 0.5].
- “alpha_slope” is the density-dependent effect on interaction strength; for each dataset, 4 values were sampled as α^ ∼ [−0.2, 0.2] within the range of [−1, 0].
- “N_opt” is the optimal neighbourhood density); for each dataset, 4 values were sampled as *N*_*o,ij*_ ∼ [0, 3] within the range of [0, 10].
- “alpha_c” is the stretching parameter; for each dataset, 4 values were sampled as α^^^ ∼ [−0.05, 0.2] within the range of [−0.5, 0].

We used the annual plant model to create the dataset (eq.7, Watkinson (1980)), with the interactions following sigmoidal functional form, embedded in the performance model used above (Ricker, 1954).

Based on each dataset, we consecutively fitted the performance model with each functional form with Bayesian inference (stan, package (The Stan Development Team, 2014), 2000 iterations and 500 warm up). Each parameter had the same prior than for the empirical application in the main manuscript: γ ∼ *N* (*means*(*fecundity*), 0), α_0_ ∼ *N* (0, 0.1), α^ ∼ *N* (−0.2, 0.2), *N*_*o,ij*_ ∼ *N* (*N*_whenFecundityismaximum_ α^^^ ∼ *N* (0, 0.1)

We see that the intrinsic performance (“lambda”) tends to be underestimated; the constant density-independent (“alpha_init”) is well-centred except with the traditional constant function, which tends to underestimate it; the density-dependent (“alpha_init”) is well-centred except for the linear function which overestimates it; the optimal density (“N_opt”) is well centred and the stretching parameter (“alpha_c” tends to be overestimate (Fig.S16). Nevertheless, zero falls within the 95% interval of each distribution. Additionally, the probability that the predefined value of the parameter falls within the 95% interval of the estimated parameter shows a common trend between functional forms (tab.S9).

**Table S9:** The probability that the predefined value of the parameter falls within the 95% interval of the estimated parameter.

| functional form | $\lambda_j$ | $\alpha_{0,ij}$ | $\hat{\alpha}_{ij}$ | $N_{0,ij}$ | $C_{ij}$ |
| --- | --- | --- | --- | --- | --- |
| Traditional constant | 67.41 | 50.47 | NA | NA | NA |
| Linear | 51.79 | 81.82 | 3.77 | 37.54 | NA |
| Exponential | 60.7 | 86.45 | 89.88 | 39.49 | 69.31 |
| Sigmoid | 58.81 | 86.48 | 92.14 | 38.49 | 68.56 |

**Figure S16:**
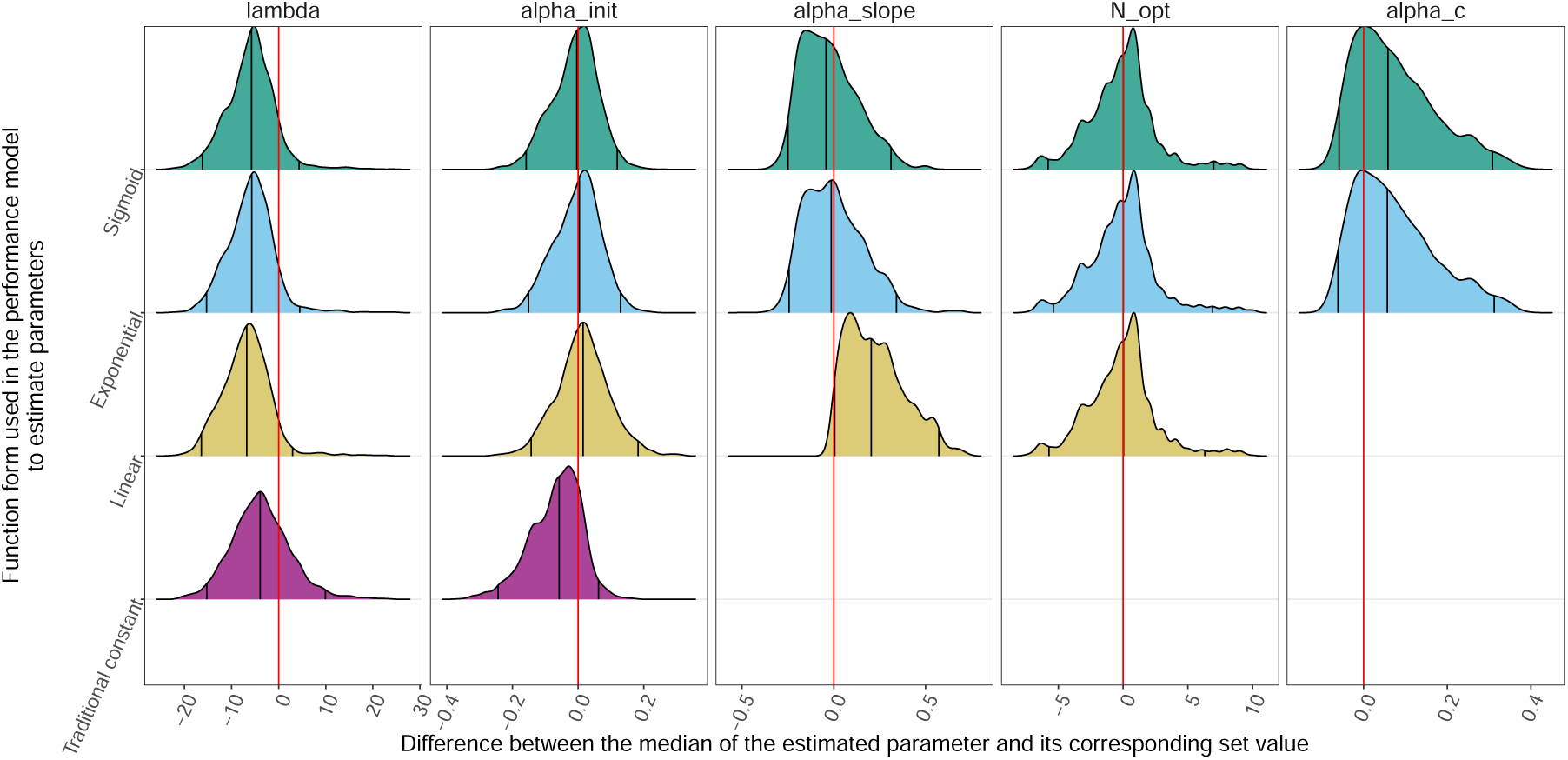
Difference between the estimated parameters and their set parameter. A positive difference means overestimation, while a negative value means underestimation. Each parameter shows the joint distributions of species *i* and species *j*. The y-axis is the functional form used to estimate the parameters in the performance model. Each column is specific to a parameter: 1-“lambda” is the intrinsic fecundity; 2-“alpha-init” is the constant density-independent effect (α_0,*ij*_ ∈ [−1, 1]); 3-“alpha_slope” is the density-dependent effect on interaction strength (e.g., α^_*ij*_ ∈ [−1, 0]); 4-“N_opt” is the optimal neighbourhood density (e.g., *N*_*o,ij*_ ∈ [0, *R*^+^]); 5-“alpha_c” is the stretching parameter (e.g., α^^^_*ij*_ ∈ [−1, 0].)

**Figure S17:**
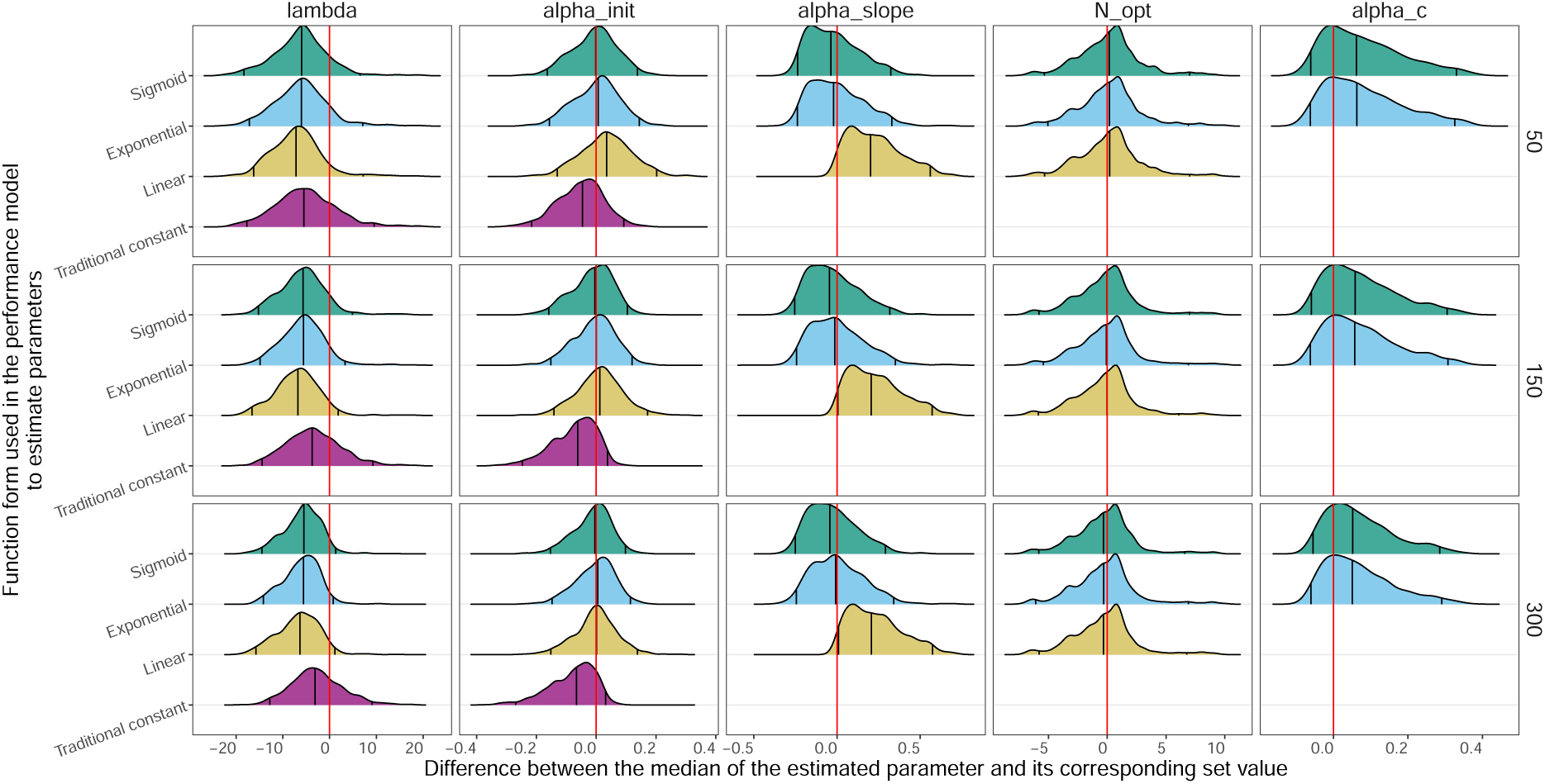
Difference between the estimated parameters and their set parameter. A positive difference means overestimation, while a negative value means underestimation. Each parameter shows the joint distributions of species *i* and species *j*. The y-axis is the functional form used to estimate the parameters in the performance model. Each row defines a number of observations (50,150, or 300) in the data used to fit the performance model. Each column is specific to a parameter: 1-“lambda” is the intrinsic fecundity; 2-“alpha-init” is the constant density-independent effect (α_0,*ij*_ ∈ [−1, 1]); 3-“alpha_slope” is the density-dependent effect on interaction strength (e.g., α^_*ij*_ ∈ [−1, 0]); 4-“N_opt” is the optimal neighbourhood density (e.g., *N*_*o,ij*_ ∈ [0, *R*^+^]); 5-“alpha_c” is the stretching parameter (e.g., α^^^_*ij*_ ∈ [−1, 0].)

**Figure S18:**
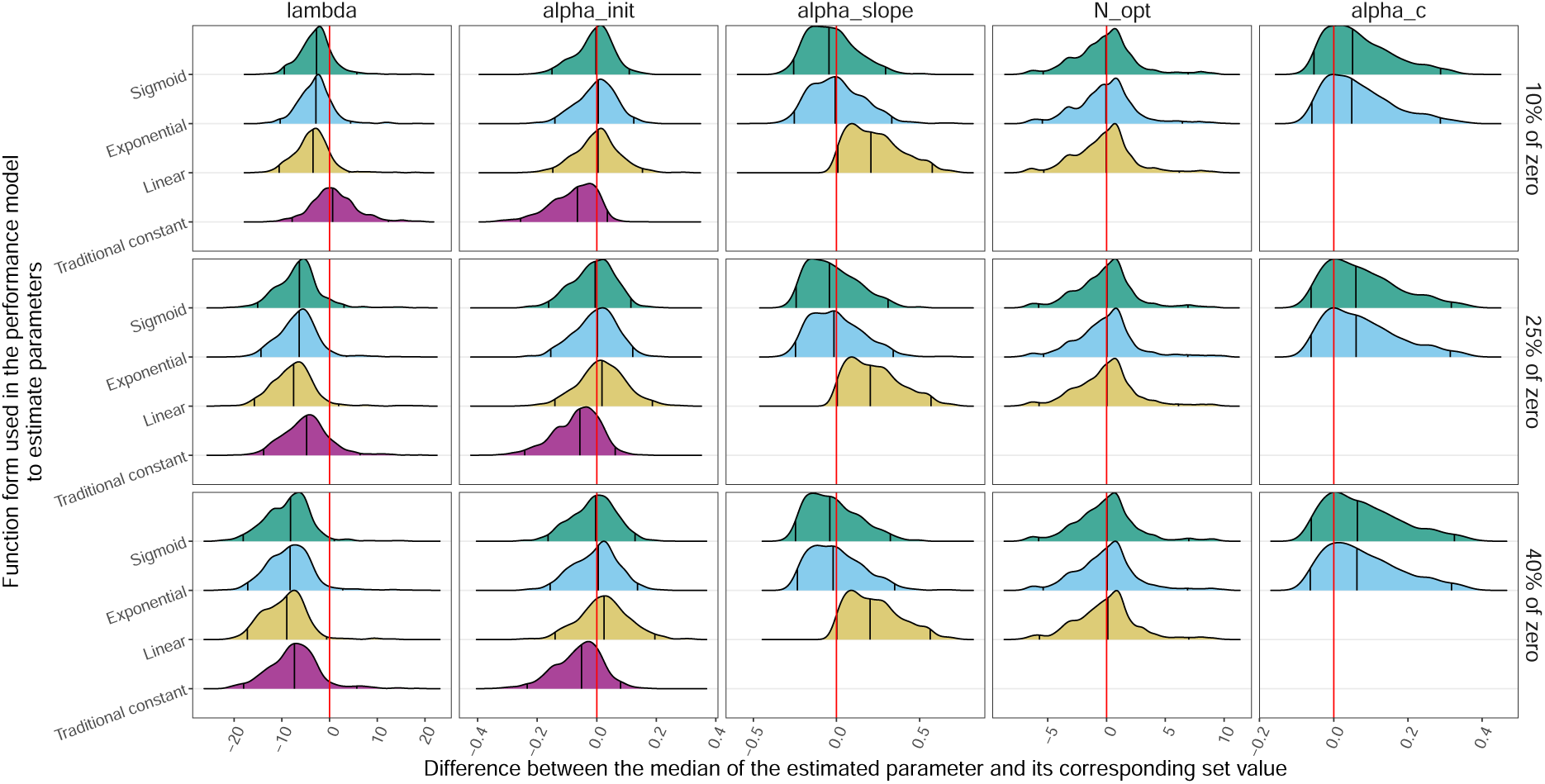
Difference between the estimated parameters and their set parameter. A positive difference means overestimation, while a negative value means underestimation. Each parameter shows the joint distributions of species *i* and species *j*. The y-axis is the functional form used to estimate the parameters in the performance model. Each row defines a percentage of zero present in the fecundity (10%,25% or 40%) used as response variable to fit the performance model. Each column is specific to a parameter: 1-“lambda” is the intrinsic fecundity; 2-“alpha-init” is the constant density-independent effect (α_0,*ij*_ ∈ [−1, 1]); 3-“alpha_slope” is the density-dependent effect on interaction strength (e.g., α^_*ij*_ ∈ [−1, 0]); 4-“N_opt” is the optimal neighbourhood density (e.g., *N*_*o,ij*_ ∈ [0, *R*^+^]); 5-“alpha_c” is the stretching parameter (e.g., α^^^_*ij*_ ∈ [−1, 0].)

Additionally, we tested the influence of two variables: (i) the number of observations (50, 150,300) and (ii) the number of performance observations equal to 0 (10, 25, or 40%). For the former variable, we see no important difference between the number of observations. For the latter variable, we see that a percentage of zero higher than 10% leads to an underestimation of the intrinsic fecundity. These two variables are mixed together when not highlighted, as in Fig.S16, where all levels of a number of observations and percentages of zero are considered together within each functional form.

## Footnotes

1 The IGR of a two-species community is theoretically computed as the ratio of the growth rates of each species when: (i) they invade at a low density, the other species at its stable density, and (ii) their respective growth rate when starting at equal density.

